# Rapid expansion and renormalization of parietal gray matter volume following associative learning

**DOI:** 10.64898/2026.04.30.721879

**Authors:** Felix Kleinschroth, Stella Villa, Antonia Lenders, Svenja Brodt, Steffen Gais, Monika Schönauer, Deniz Kumral

## Abstract

The human brain exhibits rapid structural plasticity following learning, yet its temporal dynamics and behavioral relevance remain elusive, particularly for declarative forms of learning. In this study, we tested whether T1-weighted MRI captures rapid structural reorganization following associative memory formation and whether such changes follow an expansion–renormalization trajectory. We combined three independent datasets (N = 198) to quantify gray matter volume (GMV) changes following an associative memory task using voxel-based morphometry across baseline, 2 h, and 12 h post-learning. We observed transient GMV increases in parietal, lateral occipital, and cerebellar regions at 2 h post-learning, which were no longer detectable relative to baseline by 12 h, consistent with rapid renormalization. Critically, GMV changes in left parietal cortex were associated with memory retention, such that greater maintenance of GMV was associated with better retention. In a separate, spatially distinct effect, sleep facilitated renormalization of GMV in the right parietal cortex, an effect that showed no association with memory retention. Our findings provide evidence that human gray matter undergoes hour-scale changes, reflecting dissociable memory- and sleep-related components.

## Introduction

The human brain exhibits a remarkable capacity for rapid structural reorganization in response to new experiences. While neuroplasticity has traditionally been associated with extended training periods, recent evidence suggests that measurable changes in brain tissue properties can occur within minutes to hours after learning (Villa et al., 2026). Magnetic resonance imaging (MRI) in humans offers a range of methods to study these changes (Verra et al., 2024). The most relevant are functional (f)MRI, which measures blood-oxygenation and is commonly interpreted as a measure of activity (Brodt et al., 2018; Guerra-Carrillo et al., 2014; Kourtzi et al., 2005), diffusion-weighted (DW-) MRI (Brodt et al., 2018; Friedman et al., 2025; Sagi et al., 2012), inferring tissue changes from displacement of water molecules, and T1-weighted (T1W-) MRI, which captures tissue composition and enables volumetric assessments through voxel-based morphometry (VBM; e.g. Dordevic et al., 2018; Draganski et al., 2004, 2006; Taubert et al., 2016). While an early study reported learning-induced plasticity in regions associated with visual-motor coordination following three months of juggling practice (Draganski et al., 2004), recent evidence has revised this temporal frame, providing evidence that neuroplasticity assessed by VBM can arise within minutes to hours of learning across diverse paradigms (see review: Villa et al., 2026). Rapid gray matter volume increases have been reported in a task-specific manner: visual stimulation drives changes in the visual cortex (Månsson et al., 2020; Naegel et al., 2017; Zaretskaya et al., 2023), while motor learning produces analogous gray matter changes in motor-related regions (Nierhaus et al., 2021; Olivo et al., 2022; Taubert et al., 2016). Rapid VBM-detectable plasticity following declarative or associative learning has not yet been demonstrated, despite compelling DW-MRI evidence for microstructural changes within hours of such tasks (Brodt et al., 2018).

Which dynamics characterize these rapid structural changes remains an open question. Do they persist over time, or do they reverse toward pre-learning levels? Brodt and colleagues (2018) observed mean diffusivity changes in posterior parietal cortex that persisted at 12 hours post-learning, suggesting durable microstructural modification assessed by DW-MRI. Whether T1W-MRI captures the same reorganization, and GMV changes follow a similar temporal trajectory remains unknown. One framework that speaks to the temporal dynamics of rapid structural change is the expansion-renormalization model (Wenger et al., 2017), which proposes that learning-induced structural expansions are transient, followed by

selective pruning as the system returns to homeostasis. This pattern spans multiple timescales and biological mechanisms (Lövdén et al., 2013; Wenger et al., 2017; Zatorre et al., 2012), with empirical support from rapid plasticity studies (visual: Naegel et al., 2017; motor: Taubert et al., 2010, 2016). Whether such renormalization extends to associative learning at different timescales, is detectable with T1W-MRI, and carries behavioral relevance are the questions the present study was designed to address.

To directly test whether T1W-MRI captures the same rapid structural reorganization previously observed with DW-MRI (Brodt et al., 2018; Kumral et al., 2026), we used the same associative learning paradigm and assessed whether GMV changes detected by VBM align spatially and temporally with the previously reported MD decreases, as converging evidence across imaging modalities would support the interpretation that such MRI signal changes reflect genuine structural reorganization. To address these aims, we combined three independent datasets (N = 198) with T1W-MRI acquisitions at baseline, 2 hours, and 12 hours post-learning, to characterize temporal trajectories of learning-induced volumetric changes after an associative memory task (**Fig. 1**). Notably, our dataset also included both sleep and wake intervals between scanning sessions, providing a unique opportunity to explore whether and how sleep modulates learning-induced plasticity (Bernardi et al., 2016; Brodt et al., 2023).

**Fig. 1.**
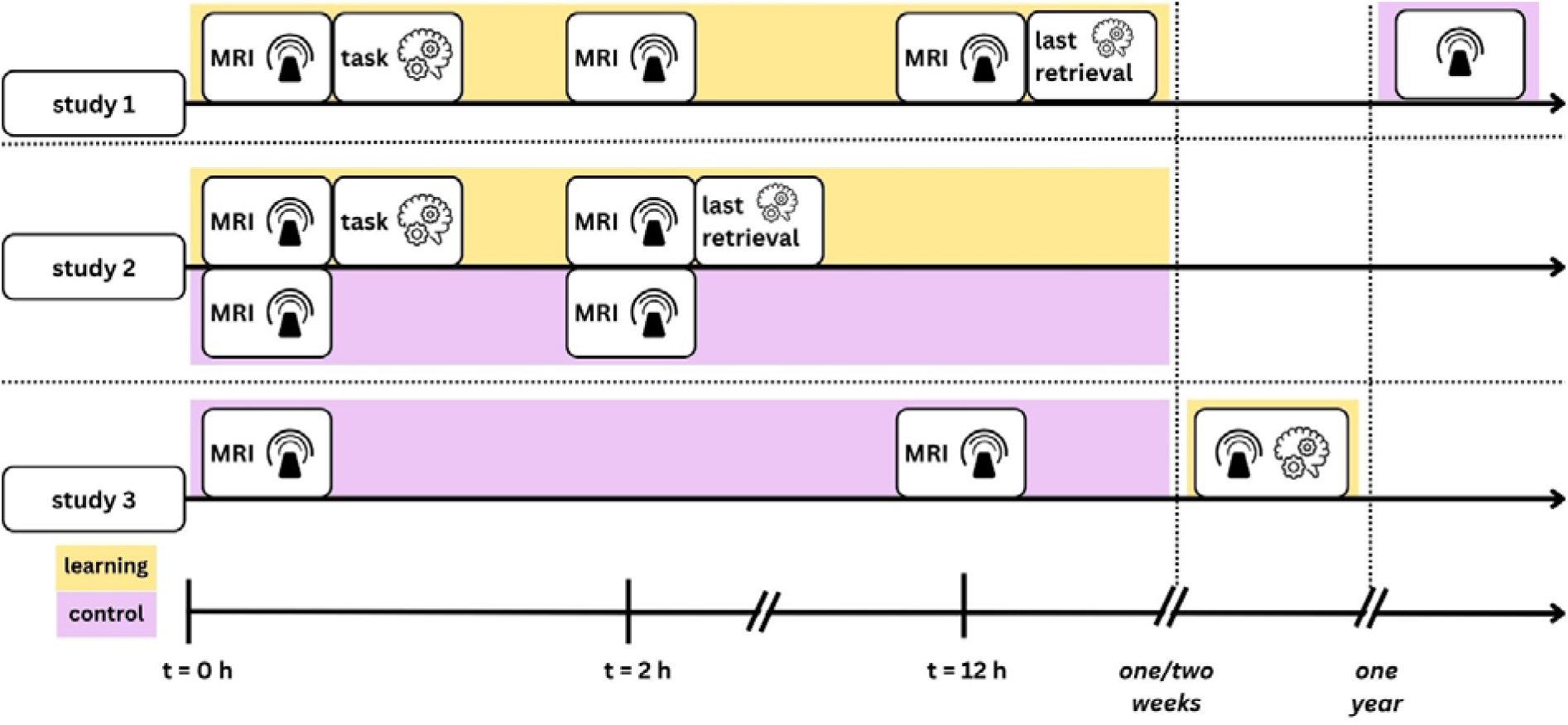
Experimental designs across three studies examining learning-induced plasticity. All studies began with a baseline MRI session (t0 = 0 h), followed by an associative memory task with 4 repetitions and subsequent post-learning MRI sessions. Participants completed a final retrieval session immediately after the last MRI scan in each study. Study 1 (N=37) used a within-subject design with MRI at t0 (baseline), t1 (≈2 h post-learning), and t2 (≈12 h post-learning), with a control session (identical procedure, no task) approximately one year later. Study 2 (N=102) used a between-subjects design with MRI at t0 and t1 only. Study 3 used a within-subject design (N=59) with MRI at t0 and t2, with control measurements one to two weeks apart. Yellow shading indicates experimental (learning) sessions; purple indicates control (no-learning) sessions. Dashed vertical lines mark critical timepoints and the slanted lines indicate time scale changes; dashed horizontal lines separate studies. t0, baseline; t1, ≈2 h post-learning; t2, ≈12 h post-learning; MRI, magnetic resonance imaging.

Our findings revealed immediate GMV increases in parietal cortex, lateral occipital cortex, and cerebellum at 2 hours post-learning, extending previous DW-MRI evidence for the rapid microstructural changes in this paradigm (Brodt et al., 2018). Early GMV increases renormalized by 12 hours in line with the expansion-renormalization framework (Wenger et al., 2017), and greater left parietal renormalization was associated with poorer memory retention. Sleep modulated this renormalization trajectory, suggesting that the sleeping brain differentially preserves learning-induced structural changes in task-relevant regions. Together, our findings establish that rapid, behaviorally relevant gray matter plasticity following associative learning is detectable with T1W-MRI, extending the expansion-renormalization framework to a cognitive domain and a sub-diurnal timescale.

## Results

### Memory performance

All three studies implemented an image-location associative memory task in which participants learned spatial locations for pairs of images (Study 1: object-object pairs; Studies 2 and 3: object and scene image pairs; see Methods section). Using a two-way mixed ANOVA with factors of repetition and study site, we found no significant interaction between repetition and study site (*F*(8, 628) = 1.72, *p* = .09, η² = .021), indicating that performance changes over time were comparable across studies. Importantly, there was also no main effect of study site (*F*(2, 157) = 0.16, *p* = .85, η² = .002), suggesting that participants performed similarly regardless of testing location. In contrast, we observed a significant main effect of the repetition on the task performance (*F*(4, 628) = 443.41, *p* < .001, η² = .74). In all three studies, participants’ performance improved from the first to the fourth retrieval phase, before declining from fourth to fifth retrieval (**Fig. 2**). Post-hoc paired t-tests with Bonferroni-correction revealed that performance increased consistently from first repetition (*M* = 15.62, *SD* = 11.58) to fourth repetition (*M* = 65.06, *SD* = 29.29, *t*(159) = - 25.41, *p* < .001), before a slight decrease from R4 to R5 (*M* = 61.69, *SD* = 30.02, *t*(159) = - 5.29, *p* < .001). These results suggest improvements in the recall of the image pair with each repetition during the initial task phase (R1-R4) and a substantial retention of the learned contents after 12 hours (R5). Because the R4–R5 interval differed across studies (∼2 h in Study 2 vs. ∼12 h in Studies 1 and 3), we additionally ran paired t-tests comparing R4 and R5 separately within each study. Memory performance was significantly lower at R5 than at R4 in Study 1 (t(36) = 5.06, *d* = -0.83, *p* < .001) and Study 2 (*t*(70) = 3.53, *d* = -0.42, *p* < .001), whereas the difference was not significant in Study 3 (*t*(51) = 1.18, *d* = -0.16, *p* = .245). Importantly, in Study 1, this reduction was driven primarily by the wake group, whereas memory performance was comparatively preserved following sleep, consistent with the sleep-dependent preservation of memory reported previously for this dataset (Brodt et al., 2023). This pattern is also consistent with the relatively preserved memory performance observed in Study 3, in which all participants underwent the sleep condition.

**Fig. 2.**
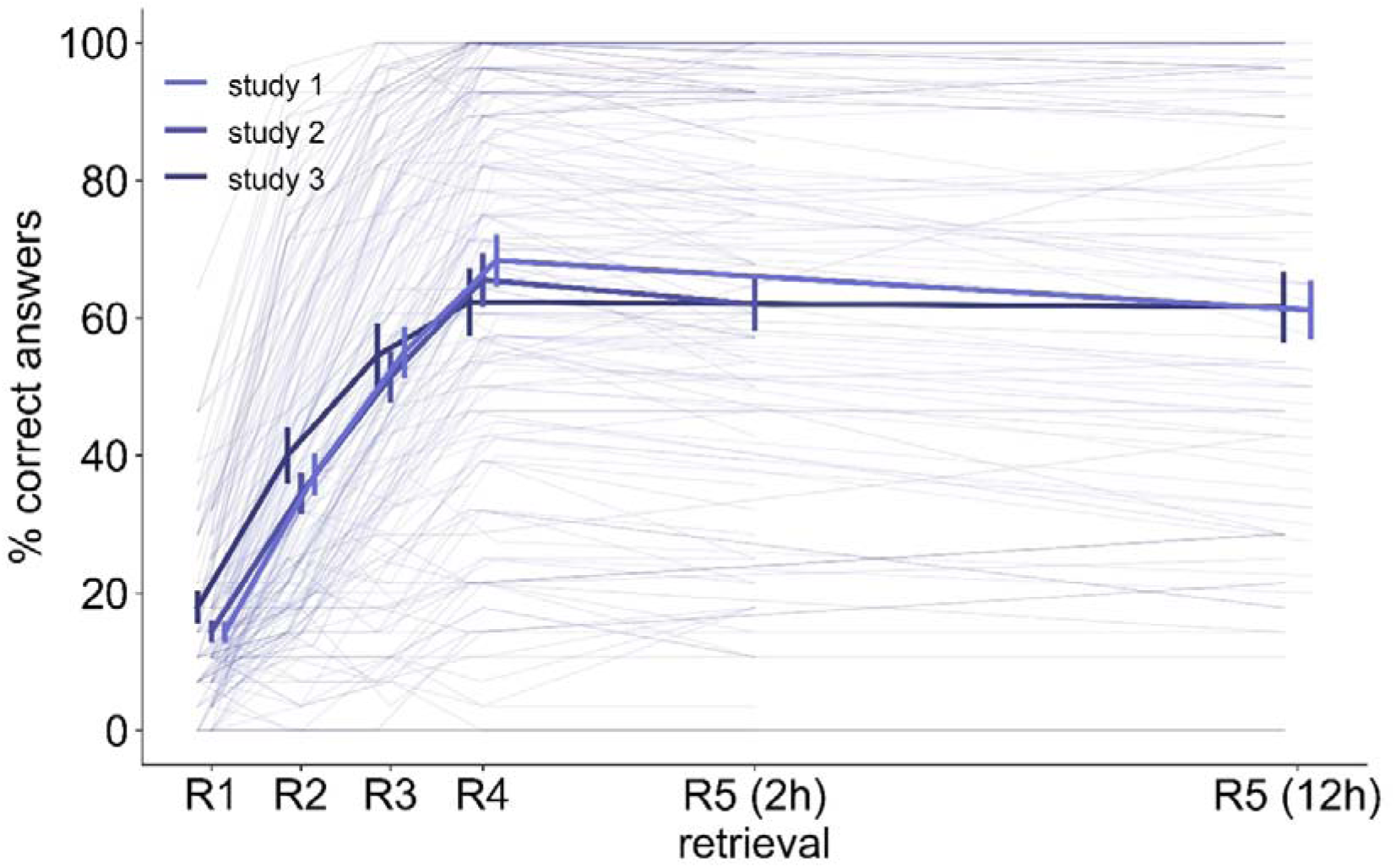
Performance across studies and repetitions. Participants’ performance on the image–location association task increased from the first (R1) to the fourth (R4) retrieval phase across all three studies. The fifth retrieval (R5) was conducted at 2 h post-learning in study 2 (labeled “R5 (2h)”) and at 12 h post-learning in studies 1 and 3 (labeled “R5 (12h)”). A mixed ANOVA confirmed a significant main effect of repetition, while post-hoc comparisons indicated significant differences between all retrieval levels (R1– R5, all *p* < 0.001). Error bars represent the standard error of the mean and individual data points are shown as transparent lines. Shade of purple color indicate the individual studies.

### GMV increases two hours after learning

Because the three studies differ in the post-learning timepoints available, different subsets of studies contribute to the following analyses. The study and sample-size contributions to each temporal comparison are summarized in **Table S3**. Using a flexible factorial model, we found an increase in GMV that was greater in the learning than in the control condition in 6 clusters spanning 5 regions, including the bilateral parietal operculum cortex (PC), lateral occipital cortex (LOC), precentral gyrus, anterior medial temporal lobe and cerebellum (**Fig. 3a**, **Table S1**; *k* > 20, *p*_uncorr_ = .001). Note that among these clusters, only the right PC survived a peak-level FWE correction (*p*_FWE_ < .05; *p*_right_PC_ = .047); we therefore selected the bilateral PC clusters for subsequent analyses (e.g., correlation with behavior). For each available participant (Study 1 + 2), the GMV change (t1-t0) was computed for the bilateral PC clusters (*n*_learning_ = 109, *n*_control_ = 62). Using linear mixed effect models (LMM), we then tested whether GMV change differed between the learning and the control conditions depending on the study site. Likelihood ratio tests showed that including the study site × condition interaction did not significantly improve the model fit for either the left (χ²(1) = 1.21, *p* = .27) or the right PC (χ²(1) = 1.88, *p* = .17), suggesting that the condition effect did not differ significantly between study sites. The null model revealed a significant main effect of study site (left: β = -0.007, *SE* = 0.001, *p* < .001; right: β = -0.010, *SE* = 0.001, *p* < .001) and the effect of condition, that the VBM analysis had detected (left: β = 0.002, *SE* = 0.001; right: β = 0.002, *SE* = 0.001). Post-hoc tests confirmed the differences between the learning and the control condition in both PC clusters (left: *t*(115.25) = -3.98, β = -0.002, *p* < .001; right: *t*(115.25) = -3.95, β = -0.002, *p* < .001). The pattern of GMV increase is also reflected in the descriptive statistics, showing greater GMV increases in the learning compared to the control condition in both clusters (left PC learning vs control: *M* = 0.00568, *SD* = 0.0050 vs. *M* = 0.00195, *SD* = 0.0051; right PC learning vs control: *M* = 0.00699, *SD* = 0.0055 vs. *M* = 0.00324, *SD* = 0.0047; **Fig. 3b, c**).

**Fig. 3.**
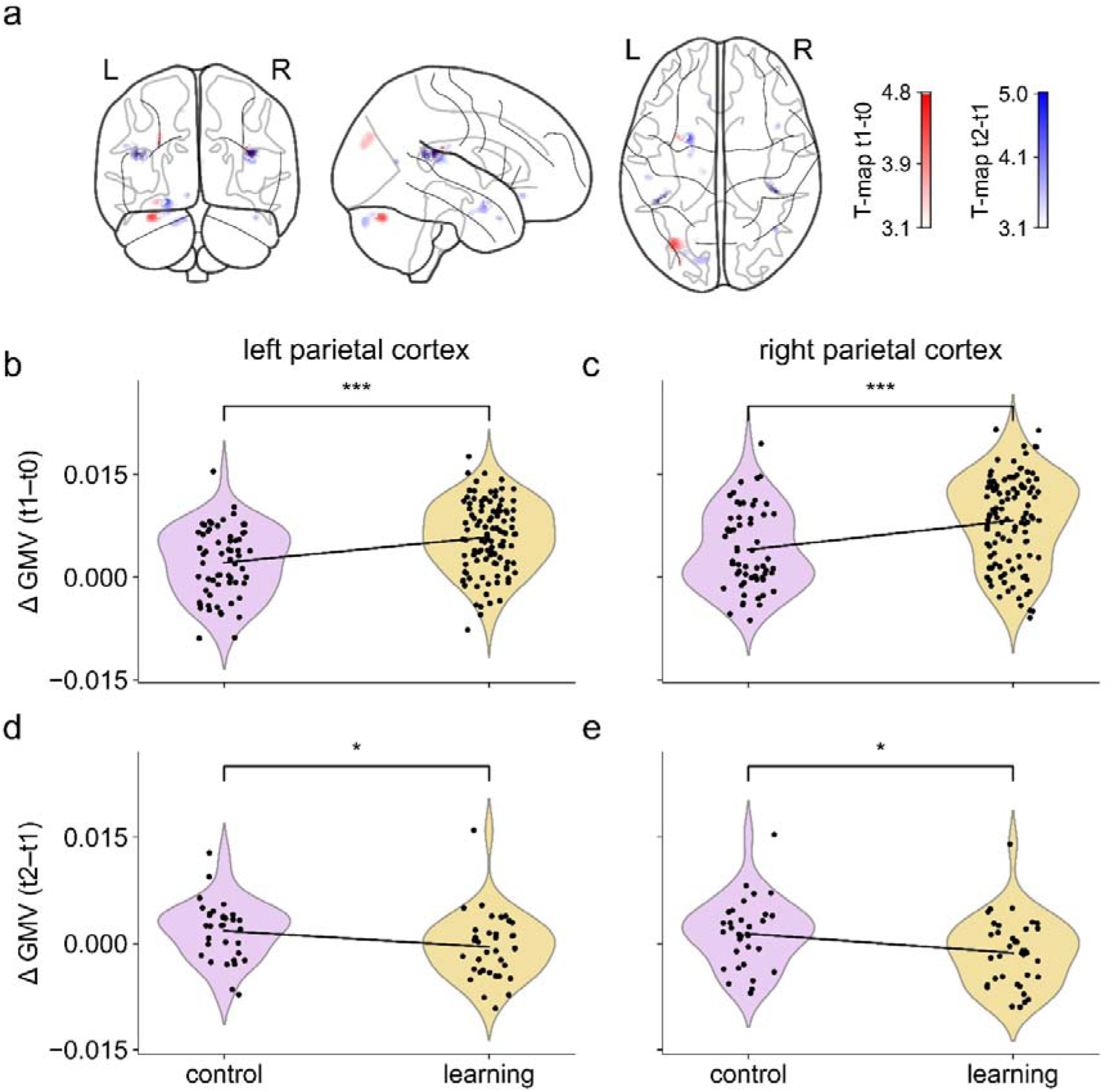
Learning-induced gray matter volume (GMV) changes at 2 h and 12 h post-learning. (**a)** Spatial maps showing significant clusters where GMV increased from baseline to 2 h post-learning in the learning relative to the control condition (red) and clusters where GMV decreased significantly from 2 h to 12 h post-learning in the learning relative to the control condition (blue). GMV change (t1-t0) in the left (*p*_uncorr_ < 0.001, *k* > 20 voxels) (**b**) and right (**c**) parietal cortex (PC) clusters for learning (yellow) and control (purple) conditions across studies 1 and 2 (*n*_learning_ = 109, *n*_control_ = 62). GMV increased more in the learning than in the control condition in both clusters. The bottom row shows GMV change (t2-t1) in the left (**d**) and right (**e**) PC clusters for study 1 only. Paired t-tests (*n*_learning_ = 32, *n*_control_ = 32) confirmed that GMV declined more in the learning than in the control condition in both, indicating renormalization specific to the learning group. Individual data points are plotted on top of the violin plots. GMV, gray matter volume; t0, baseline; t1, 2 h post-learning; t2, 12 h post-learning (Bonferroni-corrected; * p < .05, *** p < .001)

### GMV renormalization at twelve hours post-learning

To characterize GMV trajectories across the full post-learning period, we next examined whether GMV differed between baseline and 12-hour post-learning in the learning relative to the control condition (Brodt et al., 2018). As this contrast yielded no significant clusters, we subsequently examined the t2-t1 interval for GMV decreases in the learning relative to the control condition in all participants with available T1W MRI at both 2 h and 12 h after learning (study 1, n = 37) using a flexible factorial model. We identified 12 clusters across 10 regions showing significantly smaller GMV in the learning than the control condition (**Fig. 3a**, **Table S2**). These regions overlapped with those identified in the t0–t1 analysis including bilateral PC, LOC, anterior medial temporal lobe, precentral gyrus, and cerebellum with additional clusters in the temporal pole, supramarginal gyrus, and cingulate gyrus. The exact contributions of different labels to the PC clusters and anterior medial temporal cluster are listed in **Tables S6-8.** Clusters showing overlap with the early expansion that exceed our predefined extent criterion (*k* > 20 voxels) are reported in **Table S4** and corresponding MNI slices are visualized in **Fig. S4**. At peak-level FWE correction, only the right PC survived (*p*_FWE_ = .016); the right PC region also passed cluster-level correction (*k* = 8, *p* = .022, *p*_FWE_ < .05). We then assessed whether GMV change (t2-t1) differed between the learning and the control conditions using two-tailed paired t-test (*n*_learning_ = 32, *n*_control_ = 32), and observed significant differences for both left PC (*t*(31) = 2.72, *d* = 0.68, *p* = .01) and the right PC (*t*(31) = 2.47, *d* = 0.62, *p* = .02). While the average GMV increased in the control (left PC: *M* = 0.0018, *SD* = 0.0042; right PC: *M* = 0.0014, *SD* = 0.0048), there was a GMV decrease in the learning condition (left PC: *M* = - 0.0005, *SD* = 0.0046; right PC: *M* = -0.0013, *SD* = 0.0048) between 2 and 12 h after learning (**Fig. 3d, e**). A negative ΔGMV (t2-t1) indicates that GMV was greater at the 2 h time point than at the 12 h time point, reflecting a decrease in GMV between these two assessments and therefore greater structural renormalization. Values closer to zero indicate greater maintenance of the GMV increase observed at 2 h.

### The relation of GMV renormalization with memory retention

To examine associations between structural plasticity and memory, we computed Pearson correlations between GMV changes in the bilateral PC clusters from the VBM analysis at both the t1-t0 and t2-t1 intervals and memory retention, defined as the difference in performance accuracy between the fourth and fifth retrieval. While we observed no association between memory retention and short-term (within 2 h) GMV increases in PC (**Fig. S1**), memory retention was positively correlated with GMV changes between t1 and t2 in the left PC (*r* = .42, 95% *CI* [0.11, 0.65], *p* = .01, *n* = 37), indicating that greater GMV maintenance (less renormalization) in the left PC was associated with better memory retention (**Fig. 4a**). However, for the right PC we observed no association between GMV changes and memory (*r* = .15, 95% *CI* [-0.18, 0.45], *p* = .37, *n* = 37; **Fig. 4a, b**). A complementary median split analysis further confirmed this correlation: participants with above-median (left: *M* = -3.81, *SD* = 7.19, *n* = 19; right: *M* = - 6.58, *SD* = 7.23, *n* = 19) GMV maintenance between t1 (2 h) and t2 (12 h) post-learning showed significantly higher memory retention in the left PC (*t*(35) = -2.65, *d* = -0.87, *p* = .01), but not in the right PC (*t*(35) = -0.46, *d* = -0.15, *p* = .46, **Fig. 4c, d**), when compared to participants below median GMV change (left: *M* = -10.83, *SD* = 8.87, *n* = 18; right: *M* = - 7.92, *SD* = 10.19, *n* = 18).

**Fig. 4.**
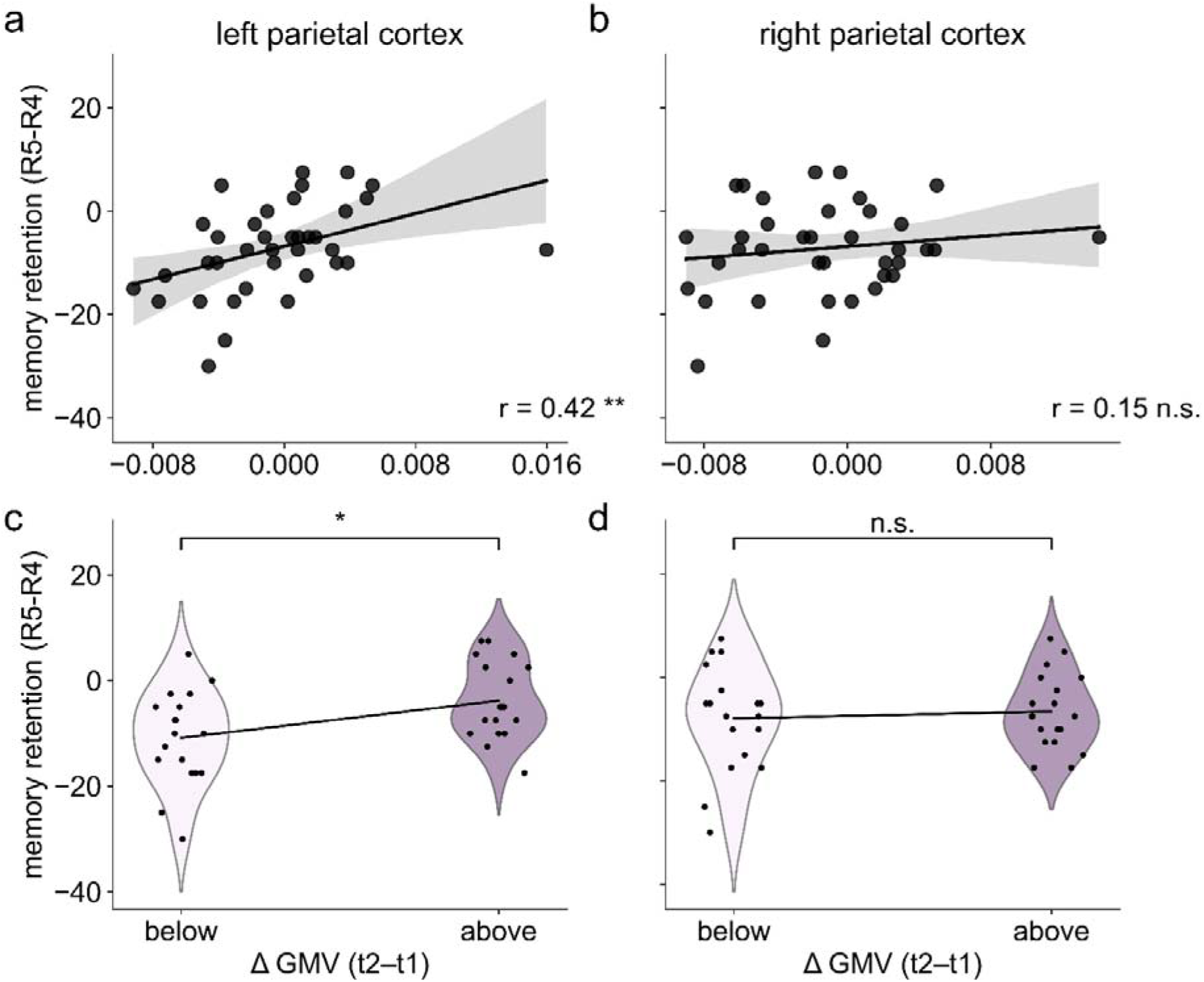
Association between GMV renormalization in parietal cortex and memory retention. Pearson correlation between GMV change (t2-t1) and memory retention in the left (**a**) and right (**b**) parietal cortex (PC) clusters, learning condition of study 1 only. A significant positive correlation was observed in the left PC but not the right PC, indicating that greater renormalization in the left PC was associated with poorer retention. Memory retention split by median GMV change (t2-t1) in the left **(c)** and right PC **(d)**, respectively. Participants with greater GMV maintenance showed higher memory retention in the left PC, but not in the right PC. Individual data points are shown over the violin plots. GMV, gray matter volume; R4, fourth retrieval; R5, fifth retrieval; PC, parietal cortex; t1, 2 h post-learning; t2, 12 h post-learning. Gray, shaded regions indicate the 95 % confidence interval (Bonferroni-corrected; * p < .05, ** p < .01, n.s. not significant).

### Sleep facilitates learning-related renormalization

Given that our dataset also included both sleep (study 1 + 3) and wake (study 1) intervals between the two post-learning sessions (t1– t2), we further examined whether the learning-related renormalization of GMV was modulated by sleep. Participants in Study 1 were randomly assigned to the evening (sleep) or morning (wake) group. The two groups did not differ significantly in chronotype, as measured by mid-sleep on workdays (MCTQ; Roenneberg et al., 2003; *t*(35) = 1.00, *d* = 0.33, *p* = .326). The comparison of sleep and wake groups across all three studies likewise revealed no significant difference in chronotype (*t*(183) = 0.77, *d* = 0.16, *p* = .44). See Supplementary **Table S5** for average measures across studies. In the clusters previously determined by the VBM analysis, the 2 × 2 mixed ANOVA with factors of sleep (sleep vs. wake; *n*_sleep_ = 17, *n*_wake_ = 15) and condition (learning vs. control, n = 32, within-subject design) revealed significant main effects of both sleep (*F*(1, 30) = 31.89, *p* < .001, η² = 0.52) and condition (*F*(1,30) = 7.66, *p* = .01, η² = 0.19) on GMV change in the left PC, with no significant interaction between the two (*F*(1, 30) = 0.01, *p* = .91, η² < 0.001). In the right PC, significant main effects of sleep (*F*(1, 30) = 16.89, *p* < .001, η² = 0.36) and condition (*F*(1, 30) = 7.33, *p* = .01, η² = 0.20) were observed, alongside a significant interaction between sleep and condition (F(1, 30) = 6.81, *p* = .014, η² = 0.19). Consistent with prior evidence for circadian modulation of brain structure (Trefler et al., 2016), GMV increased in the sleep group (left PC: *M* = 0.0029, *SD* = 0.0041, right PC: *M* = 0.0017, *SD* = 0.0051) relative to the wake group (left PC: *M* = -0.0029, *SD* = 0.0033, right PC: *M* = -0.0020, *SD* = 0.0030) between t0 and t2 in both PC clusters. To further characterize the significant sleep × condition interaction, we implemented post-hoc comparisons between the learning and the control condition. Our analysis revealed significant differences between the learning and the control condition in the right PC only in the sleep group (*t*(16) = 3.54, *d* = 0.86, *p* = .01), with greater GMV renormalization observed in the learning (*M* = -0.0006, *SD* = 0.0060) compared to the control condition (*M* = 0.0047, *SD* = 0.0031). This pattern was absent in the wake group, suggesting that sleep preferentially amplifies renormalization of learning-induced GMV changes in the right PC. All remaining comparisons were non-significant (**Fig. 5**; right PC, wake: *t*(15) = 0.053, *d* = 0.01, *p* = 1.0; left PC, sleep: *t*(16) = 1.84, *d* = 0.45, *p* = 0.34; left PC, wake: *t*(15) = 2.01, *d* = 0.52, *p* = .26, Bonferroni-corrected for 4 comparisons).

**Fig. 5.**
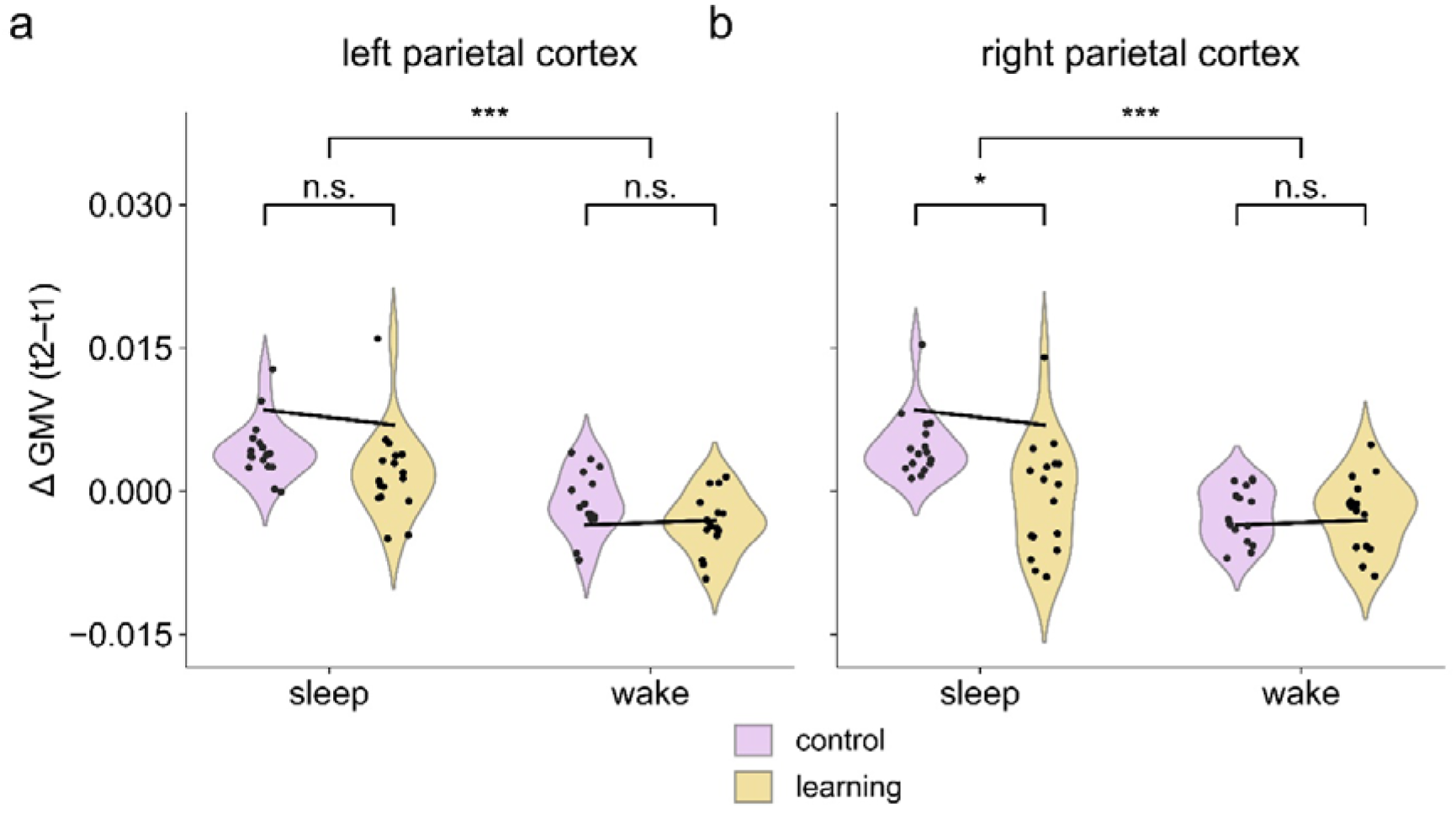
Sleep modulates GMV renormalization in the right parietal cortex. GMV change between 2 h and 12 h post-learning (t2-t1) in the left (**a**) and right (**b**) parietal operculum cortex (PC) by sleep condition (study 1 only, within-subject design; sleep: *n* = 17; wake: *n* = 15). A 2 × 2 mixed ANOVA confirmed a significant main effect of sleep and condition in the left and right PC, as well as an interaction between the two in the right PC. Post-hoc comparisons revealed greater renormalization in the learning than in the control condition within the sleep group only in the right PC (*t*(16) = 3.54, *d* = 0.86, *p* = .01), with all remaining comparisons non-significant (all *p* > .26, Bonferroni-corrected). GMV, gray matter volume; t0, baseline; t2, 12 h post-learning (Bonferroni-corrected; * p < .05, *** p < .001, n.s. not significant).

When the same analysis was applied to GMV change between baseline and the 12-hour post-learning scan (t2-t0, N=91, within-subject design), only the main effect of sleep remained significant in the bilateral PC, with no significant effect of condition or sleep × condition interaction (**Fig. S2,** all *p* > .05, Bonferroni-corrected), indicating that the learning-related structural effects (i.e., renormalization) are transient and specific to the 2- to 12-hour post-learning interval.

We lastly explored whether the association between GMV renormalization (t2-t1) and memory retention differed between the sleep and wake groups by computing Pearson correlations separately for wake and sleep groups. No significant associations were observed in either group or cluster (**Fig. S3,** all *p* > .05), likely reflecting insufficient power in the resulting subgroups. Notably, the correlations in the left PC were positive in both the sleep (*r* = .22) and wake (*r* = .32) groups, directionally consistent with the whole-sample effect, suggesting that the GMV–retention relationship in the left PC may persist across consolidation conditions but cannot be evaluated reliably at this sample size.

### Control Analysis

To assess the anatomical specificity of the observed effects, we implemented a control analysis using a task-unrelated brain region. GMV was extracted separately from the left and right juxtapositional lobule cortex (JLC) using the Harvard-Oxford cortical structural atlas (Desikan et al., 2006), and the same statistical analyses applied to the bilateral parietal operculum were repeated using these control-regions. Neither left nor right JLC GMV showed a significant effect of learning or sleep, and neither was significantly associated with memory retention (**Figs. S5-7**). Thus, the absence of corresponding effects in the JLC supports the anatomical specificity of the effects observed in the bilateral parietal operculum.

## Discussion

The present study examined whether T1W-MRI is sensitive to rapid, learning-induced structural reorganization following an associative memory task. Combining three independent datasets (total N = 198, with study-specific subsamples contributing to individual analyses; see **Table S3**), we observed transient increases in GMV across parietal, lateral occipital, and cerebellar regions at two hours post-learning. However, these early GMV increases were no longer detectable relative to baseline 12 hours post-learning, indicating renormalization. Changes in left parietal GMV were further linked to how well participants retained the memory, such that greater renormalization was associated with poorer retention. Notably, sleep facilitated renormalization of learning-induced GMV increases in parietal regions, suggesting that the post learning sleep may contribute to the normalization of structural changes in task-relevant areas.

The early GMV increases observed here are consistent with a growing body of evidence demonstrating that learning-induced structural changes can be detected within minutes to hours using VBM (see Villa et al., 2026). The review further indicates that rapid consolidation effects following cognitive learning tasks have been reported almost exclusively using DW-MRI, with such paradigms remaining underrepresented in the VBM literature. Rapid GMV increases have been reported in task-relevant cortical areas following visual stimulation (Naegel et al., 2017; Zaretskaya et al., 2023), balance training (Taubert et al., 2016), and motor-imagery practice using brain-computer-interface (Nierhaus et al., 2021). The temporal profile of the observed GMV changes also relates to the broader dynamics of memory consolidation. Consolidation of newly formed memories is further associated with dynamic changes in hippocampal–neocortical interactions during the hours following learning. Shortly after encoding, hippocampal coupling with widespread cortical networks increases (Folvik et al., 2023), but this increase returns towards baseline by 12 hours, when previously learned items elicit stronger cortical BOLD responses than novel items (Himmer et al., 2019). Over 24 hours, spanning a night of sleep, hippocampal activity and hippocampal–neocortical connectivity further decrease, whereas activity and connectivity within neocortical regions increase (Takashima et al., 2009), suggesting progressive reorganization of memory-related networks during consolidation. By comparison, structural evidence spanning a similar post-learning window remains limited. To our knowledge, no VBM study using an associative memory paradigm has characterized structural changes from the hours following learning to the next day, leaving this intermediate post-learning window largely unexplored. The GMV changes observed here therefore extend the predominantly functional literature on post-learning consolidation by providing evidence of structural reorganization across this early post-learning period. Although structural measures cannot directly establish the functional mechanisms underlying memory consolidation, our findings are consistent with the broader view that newly encoded associations undergo continued neural reorganization beyond the initial encoding period.

In our study, bilateral PC emerged as the most affected region, showing both the early GMV increase and the subsequent renormalization. This finding is notable because the parietal operculum encompasses secondary somatosensory cortex and adjacent multisensory convergence zones, integrating tactile, visual, and proprioceptive signals to form visuospatial body maps, serving as a gateway between primary sensory cortex and higher associative areas (Burton et al., 2008; Jung et al., 2009; Sirigu & Desmurget, 2021). Because the image–location task requires binding visual object identity to allocentric spatial position (Postma et al., 2008), learning may recruit parietal representations that extend beyond purely visual processing. This is consistent with evidence that the parietal operculum encodes object properties relevant to subsequent behavior in tactile paradigms (Maule et al., 2015), suggesting that this region represents object-related information in a modality-independent way. Repeated visuospatial associations may therefore also modify parietal regions that support this kind of multisensory representation, rather than acting on visual processing alone. Our cluster also extended into the planum temporale and, to a lesser extent, the inferior parietal lobule and supramarginal gyrus, encompassing a transitional region known as the temporoparietal junction (TPJ). Although the TPJ does not have clearly defined anatomical boundaries (Bzdok et al., 2013; Doricchi, 2022), it has been proposed as a site of convergence across processing streams (Carter & Huettel, 2013; Patel et al., 2019) and of attentional orienting during episodic retrieval (Ciaramelli et al., 2008; Corbetta & Shulman, 2002). This extension may reflect engagement of integrative and attentional mechanisms supporting the binding and retrieval of learned visuospatial associations, complementing the multisensory processing supported by the parietal operculum itself. While these functional findings are not to be understood as evidence of structural change in these areas, they provide a link through the activity dependent biological consolidation mechanisms discussed later on. Together, these results suggest the observed GMV changes reflect structural adaptation to the perceptual, spatial, multisensory, and attentional demands of learning, rather than a direct structural trace of the memories themselves.

Beyond the parietal regions, rapid GMV increases were also observed in the LOC, anterior medial temporal lobe, precentral gyrus, and cerebellum. The LOC is the primary cortical region for visual object recognition (Doniger et al., 2000; Grill-Spector et al., 1998, 1999), and its recruitment during associative encoding is well documented: LOC and fusiform activity at encoding predict subsequent associative memory performance, and functional coupling between these occipitotemporal regions and the medial temporal lobe supports successful item-context binding (Bencze et al., 2024). Regarding the anterior medial temporal cluster, we also observed partial spatial overlap between the initial GMV increase and the subsequent decrease, raising the possibility that an expansion–renormalization process occurs within partially overlapping anatomical territory. This cluster spans a transition zone within the anterior medial temporal region, encompassing structures such as the amygdala, parahippocampal, and olfactory cortices, with the larger renormalization cluster also extending into the hippocampus. The medial temporal lobe has a well-established role in memory formation, with the hippocampus considered a key site for the formation and consolidation of newly acquired memories (Frankland & Bontempi, 2005; McGaugh, 2000). Within this system, the hippocampus appears especially important for associative and object–location binding, while parahippocampal regions have similarly been linked to encoding object–location associations (Fromm et al., 2026; Gillis et al., 2016; Hampstead et al., 2011; Ross & Slotnick, 2008). The amygdala, too, can contribute to associative memory, including memory for object–context associations, although its role is most firmly established in modulating memory formation under emotionally arousing conditions (Barsegyan et al., 2014; McGaugh, 2004; Phelps, 2004; Roozendaal et al., 2008). Given this anatomical heterogeneity, we interpret the finding as involving the broader anterior medial temporal region rather than attributing it to a single structure. Thus, the observed structural changes are consistent with learning-related involvement of anterior medial temporal circuitry, but the anatomical heterogeneity of the clusters precludes assigning the effects to a specific medial temporal structure.

The cerebellar GMV changes observed here are consistent with the cerebellum’s well-established contributions beyond motor control (Stoodley & Schmahmann, 2009), encompassing attention regulation (Allen et al., 1997; Daniels et al., 2003), working memory (Cairo et al., 2004; Chen & Desmond, 2005), and decision making (Blackwood et al., 2004; Harrington et al., 2004). Note that the cerebellum generates predictions about upcoming sensory states and updates internal models in response to prediction errors, a domain-general computation that extends to cognitive as well as motor learning (Kostadinov et al., 2019; Sokolov et al., 2017; Wolpert et al., 1998). These predictive computations are plausibly recruited during the image-location task, where the cerebellum may track statistical regularities of object-location pairings and signal violations of expectation. Computational models further suggest that cerebellar-driven cortical dynamics facilitate rapid task learning, with task-relevant information gradually transferred from the cerebellum to the cortex during consolidation (Boven et al., 2023; Boven & Cerminara, 2023), consistent with the transient nature of the cerebellar GMV changes observed here. Taken together, these findings may reflect rapid structural remodeling across multiple components of a distributed learning network. The concurrent changes observed across medial temporal, parietal, and cerebellar regions suggest that learning-related plasticity may emerge across interconnected regions rather than reflecting a simple transfer of information from one region to another.

A central finding of the present study is that the early GMV increases observed at two hours post-learning were no longer detectable 12 hours later. Rather, when GMV change between the 2-hour and 12-hour timepoints was examined, the learning condition exhibited a reduction in GMV relative to the control condition across regions overlapping with those that had shown initial expansion. This pattern of initial GMV increase followed by a decrease toward baseline is consistent with the expansion-renormalization model (Wenger et al., 2017), which proposes that learning transiently recruits neural resources including dendritic spine formation, glial swelling, and associated water redistribution, before a selective stabilization process prunes non-optimal elements as the system approaches homeostasis (Wenger et al., 2017). Previous evidence for renormalization has been reported across multiple timescales: visual cortex GMV returned to baseline within 60 minutes of cessation of flickering checkerboard stimulation (Naegel et al., 2017), and cortical thickness increases in M1 after one hour of balance training were absent two weeks later (Taubert et al., 2010, 2016). The present study demonstrates that renormalization can occur across a timescale of hours following a single cognitive learning session, bridging rapid sensory/motor effects with higher-order associative memory processes. Critically, individual differences in the degree of renormalization were behaviorally relevant: greater maintenance of learning-induced GMV in the left PC between 2 and 12 hours post-learning was associated with better memory retention, whereas greater renormalization was associated with poorer retention. Notably, no such association was observed for the initial GMV increase at 2 hours, indicating that it is not the magnitude of the initial structural expansion *per se* that predicts memory, but rather how much of it is preserved through the consolidation window. This dissociation is consistent with the view that renormalization is a selective rather than uniform process, with the degree of structural preservation reflecting the strength or efficiency of initial encoding and establishing rapid plasticity as behaviorally relevant, rather than merely transient remodeling without behavioral consequence.

The cellular basis of rapid GMV changes detected by VBM remains an open question; however, synaptogenesis and glial activity, in particular astrocytic remodeling, are plausible candidates at this timescale (Zatorre et al., 2012). Structural vascular changes associated with neurogenesis (Pereira et al., 2007) persist over weeks (Rhyu et al., 2010), and neurogenesis itself is largely restricted to the dentate gyrus and subventricular zone (Eriksson et al., 1998; Kempermann et al., 2015; Kuhn et al., 2001; Ponti et al., 2013), with newly born cells requiring weeks to reach maturity (Jiang et al., 2016; Lanjewar & Sloan, 2021; Westerlund et al., 2003). These mechanisms are therefore unlikely to contribute meaningfully to structural changes that arise and renormalize within hours. Dendritic spine density increases have been directly coupled to VBM-detectable GMV changes in rodents (Keifer et al., 2015), indicating synaptogenesis as a plausible structural correlate. Microglial pruning of excess spines represents a plausible cellular mechanism for the renormalization observed here (Liu et al., 2021; Pan & Monje, 2020; Tremblay et al., 2010). Supporting this idea, depletion of microglia impairs learning in rodents (Parkhurst et al., 2013), indicating that microglial activity is functionally integral to consolidation. Beyond dendritic spines and microglia, astrocytes have been proposed as possible contributors to MRI-detectable plasticity. At a ratio of approximately 1.4 astrocytes per neuron, they represent a substantial fraction of brain tissue volume and undergo rapid, activity-dependent morphological changes within minutes in response to learning-related activity (Theodosis et al., 2008), directly altering local tissue geometry and extracellular water distribution (Degl’Innocenti & Dell’Anno, 2023; Schmidt et al., 2021; Theodosis et al., 2008). Crucially, Bernardinelli and colleagues (2014) demonstrated that the astrocytic process predicts spine plasticity, suggesting that both biological mechanisms rely on each other and may jointly contribute to the T1W signal changes (Schmidt et al., 2021). Note that hemodynamic contributions to VBM changes cannot be excluded entirely, given that vasodilation can produce apparent GMV overestimates detectable by VBM (Tardif et al., 2017). Short-lived hemodynamic responses to increased energy demand (Lewis et al., 2018) are unlikely to account for our results, as these effects resolve within seconds to minutes of task cessation. Consistent with this, the pattern of early GMV expansion followed by renormalization observed here was specific to the learning condition. The present findings are thus best interpreted as reflecting a combination of transient astrocytic remodeling and synaptogenesis (Zatorre et al., 2012).

The sleep effect observed here reflects two concurrent processes. Sleep was associated with a general GMV increase irrespective of condition, consistent with established time-of-day variation in brain volume (Nakamura et al., 2015; Trefler et al., 2016). Furthermore, extending prior evidence that recovery sleep reverses training-induced diffusivity changes in cortical gray matter (Bernardi et al., 2016), the present findings demonstrate that sleep also promotes renormalization of learning-induced GMV changes, with right PC showing greater GMV reduction in the learning relative to the control condition following sleep but not wakefulness. Our findings align with accumulating evidence that sleep modulates learning-induced plasticity and are consistent with the synaptic homeostasis hypothesis (Tononi & Cirelli, 2006). This framework proposes that experience during wakefulness causes synaptic potentiation and associated cellular changes such as astrocytic swelling and reduced extracellular space (Degl’Innocenti & Dell’Anno, 2023; Keifer et al., 2015; Schmidt et al., 2021; Theodosis et al., 2008), which sleep subsequently reverses to maintain the brain’s overall capacity for plasticity. Together, our results reveal dissociable relationships underlying hour-scale structural reorganization after associative learning: memory performance was associated with GMV maintenance in the left PC, and structural renormalization was associated with sleep in the right PC, with no evidence that sleep modulated the relationship between GMV maintenance and memory retention. Rather than reflecting a single sleep-dependent mechanism, these findings suggest complementary roles for structural maintenance and sleep-associated renormalization. Superior retention may reflect the persistence of sufficiently robust learning-related GMV changes despite subsequent renormalization, supporting successful consolidation. This interpretation does not imply, however, that the remaining GMV itself constitutes the memory trace.

A key strength of the present study is the combination of three independent datasets (N = 198) with a shared experimental design, providing substantially greater statistical power than any individual study in the rapid structural plasticity literature. The longitudinal VBM pipeline with ComBat harmonization addressed between-site variability, and the inclusion of matched control sessions with comparable scan timing and no learning task was essential for distinguishing learning-induced from time-of-day or repeated-scanning effects (Broessner et al., 2021). However, several limitations warrant consideration. Despite harmonization, inter-study differences cannot be fully eliminated and may contribute to residual heterogeneity in learning-related effects. ComBat reduces systematic scanner-related variation, but it cannot fully account for differences in acquisition protocols, experimental procedures, or temporal sampling equivalence across studies. Moreover, harmonization itself relies on modelling assumptions and therefore cannot exclude more complex scanner-by-condition or scanner-by-time interactions. These considerations warrant caution when interpreting effects that depend on comparisons across studies, although the primary learning-versus-control comparisons were performed within studies under matched acquisition conditions. Furthermore, despite the relatively large combined sample, detected clusters did not survive cluster-level FWE correction, likely reflecting the small effect sizes expected for rapid structural changes (Villa et al., 2026). Notably, not all studies included data at every interval. Analyses of the renormalization and sleep were restricted to Study 1, the only dataset including both the 2-hour and 12-hour timepoints and the sleep–wake manipulation, with the sleep comparison further subdividing this sample into small subgroups. Future studies with larger or more evenly distributed samples could benefit from using a discovery dataset to identify volumetric changes and an independent confirmatory dataset to examine their associations with sleep or behavior, thereby ensuring complete independence between the analyses.

The parietal operculum clusters identified here are anatomically distinct from the posterior parietal cortex where Brodt et al. (2018) reported MD decreases, indicating that the two modalities do not provide spatially convergent evidence but likely index different aspects of the broader parietal reorganization that follows associative encoding. The two modalities (DW-MRI and T1W-MRI) diverged temporally as well: while MD changes persisted to the 12-hour post-learning scan (Brodt et al., 2018), GMV increases had renormalized by that timepoint. These discrepancies likely reflect genuine differences in what each modality captures rather than inconsistent underlying biology (Zatorre et al., 2012). Nevertheless, the repeated cluster pattern across both time windows and its alignment with the expansion-renormalization framework support the interpretation that the present findings reflect genuine learning-related structural reorganization on a short-time scale. Future studies acquiring T1W-MRI and DW-MRI alongside complementary measures such as arterial spin labeling (Detre et al., 2012) within the same participants will be needed to disentangle the biological contributions to each signal and establish whether the two modalities index complementary or overlapping aspects of the same underlying process.

Using three independent datasets, we demonstrate that T1W-MRI is sensitive to rapid, transient GMV increases following associative learning that renormalize within 12 hours. Renormalization in left PC was associated with memory retention, while sleep-related modulation was observed in right PC, suggesting that sleep acts on structural changes specific to learning rather than reversing all wakefulness related changes equally, and that these processes are related but anatomically distinct. These findings extend the expansion-renormalization framework to a cognitive domain and sub-diurnal timescale and establish rapid parietal plasticity as a robust, behaviorally relevant correlate of memory consolidation detectable with T1W-MRI.

## Methods

### Study Design

Data from three independent studies with highly comparable experimental designs were used. In each study, participants first completed a baseline MRI session, followed by an image-location learning task and subsequently underwent a post-learning MRI session (**Fig. 1**). However, the delay between the learning task and the post-learning MRI varied across studies: study 1 included both intervals (Brodt et al., 2018, 2023), whereas study 2 used a delay of ≈2 h and study 3 a delay of ≈12 hours. Additionally, studies 1 and 3 employed within-subject designs, whereas study 2 used a between-subjects design (**Fig. 1**). Study 1 was conducted at the Max-Planck-Institute for Biological Cybernetics, Tübingen, and studies 2 and 3 were conducted at the Department of Radiology, University Medical Center Freiburg, Germany. Written informed consent was obtained from all participants, and all studies were conducted in accordance with the Declaration of Helsinki and were approved by the local ethics committee (approval numbers: 652/2015BO2, 21-1553, 23-1380).

### Participants

Participants were recruited via flyers, posters, online advertisements and, in case of studies 2 and 3 the university participant pool (SONA). Eligibility was assessed through a structured telephone screening. Inclusion criteria were age (18 to 35 years), at least B2-level proficiency in German, no history of neurological or psychological disorders, no chronic pain or sleep disorders, and no contraindications for MRI. In study 1, each subject received compensation of €12/h (Brodt et al., 2018), while in other studies participants were compensated at a rate of €10/h for behavioral, €15/h for neuroimaging, and €60/h for sleep sessions (only study 3).

In total, 198 subjects (*M* = 23.8, *SD* = 3.2 years; 106 females) were included in our analysis. Study 1 included 39 individuals who completed the learning protocol (Brodt et al., 2018). Two individuals were excluded due to insufficient image preprocessing quality, assessed by contrast-to-noise ratio (*n* = 2), resulting in a final sample of 37 participants (*M* = 24.1, *SD* = 2.7 years; 21 females), of which 32 participants returned one year later for the control (no-learning) condition. For study 2 (Kumral et al., 2026), in total 125 eligible participants were recruited, with 84 assigned to the learning condition and 41 to the control condition. Participants were excluded due to incidental findings (*n* = 2), falling asleep during data acquisition (*n* = 1), or withdrawing due to discomfort in MRI (*n* = 4), resulting in an initial sample of 118 individuals. Additional exclusions were made due to incomplete datasets (*n* = 12), MRI related artifacts (n = 3), or insufficient image preprocessing quality (*n* = 1). The final sample of study 2 included a total of 102 participants, with 72 in the learning group (*M* = 23.2, *SD* = 2.9 years; 38 females) and 30 in the control group (*M* = 24.3, *SD* = 3.0 years; 15 females). Lastly, a total of 82 participants were recruited for study 3. Of these, 68 participants completed both the learning and the control condition. After exclusions due to incomplete datasets (*n* = 8), or insufficient image preprocessing quality (*n* = 1), 59 individuals (*M* = 24.3, *SD* = 3.9 years; 32 females) remained. For a detailed overview of the selection process, see **Fig. 6**.

**Fig. 6.**
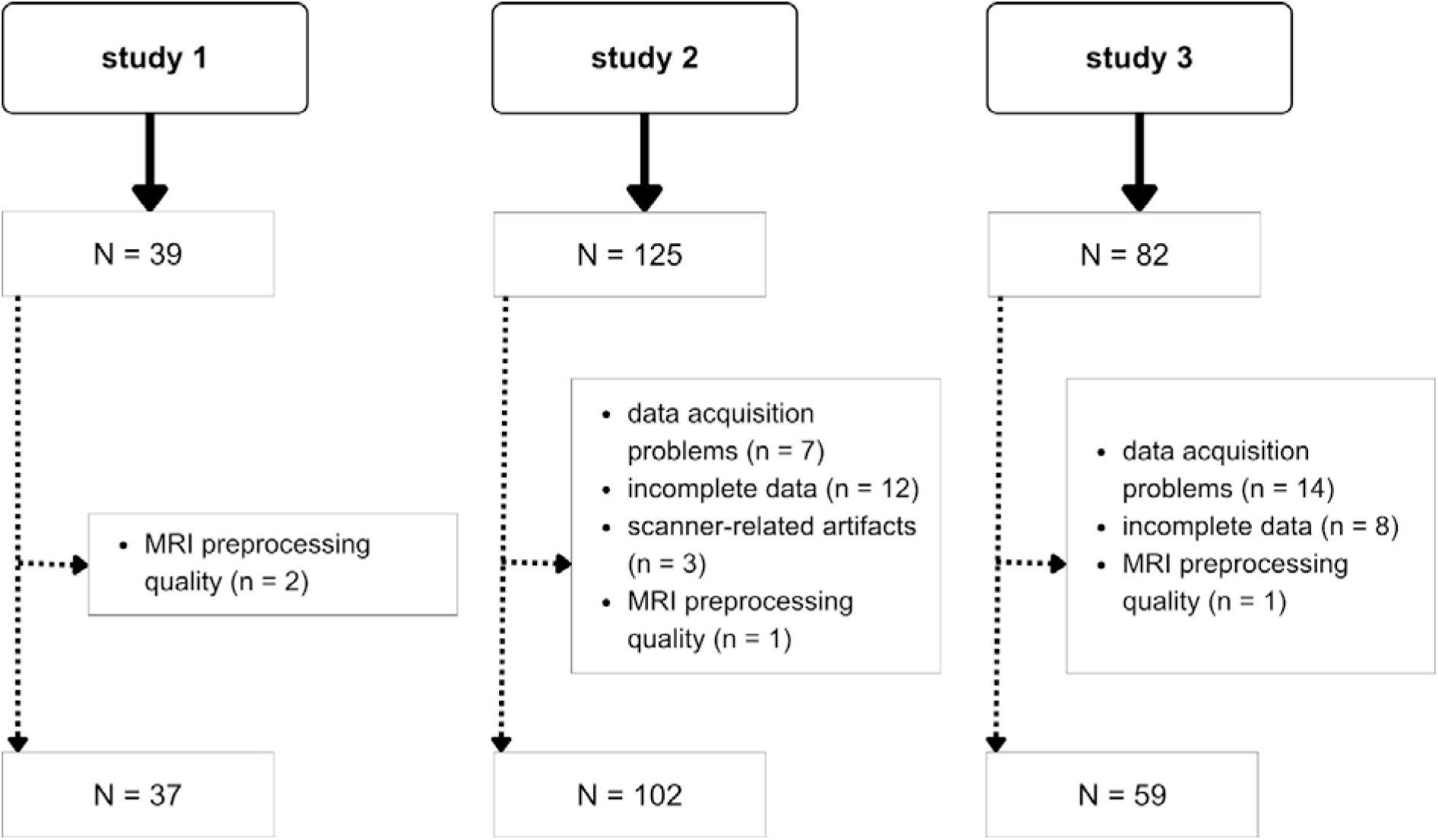
Participant selection flowchart. Of 246 screened participants, 198 were included in the final analysis. Exclusions comprised: 21 due to data acquisition problems (study 2: *n* = 7; study 3: *n* = 14), 20 due to incomplete datasets (study 2: *n* = 12; study 3: *n* = 8), 4 due to insufficient image preprocessing quality, defined as at least one image with a z-scored noise-to-contrast ratio (*NCR*) > 3, assessed within each study prior to preprocessing (study 1: *n* = 2; study 2: *n* = 1; study 3: *n* = 1), and 3 due to scanner-related artefacts identified during visual inspection (study 2: *n* = 3).

### Image-location task

In all studies, participants completed an image–location association task consisting of four encoding-retrieval blocks. Before the task started, participants received detailed instructions and completed a short tutorial, to familiarize themselves with the procedure and controls. In study 1 and study 2, participants completed the task while still in the MRI after the baseline scans; in study 3 it was done outside the scanner.

#### Study 1

Forty images were randomly placed on an 8-by-5 grid. These images depicted items from three classes: inanimate objects (*n* = 20), animals (*n* = 10), and fruits and vegetables (*n* = 10). Each image was part of two different association pairs as either the first or the second image of the pair, resulting in a total of 40 distinct association pairs. Notably, association pairs always featured one inanimate object and one object from one of the other two classes. During encoding, participants were presented with a random sequence of all association pairs. Each image was displayed at its grid location for a duration of 2 seconds, with a delay of 0.4-0.6 seconds between images belonging to the same pair and a delay of 1.55-1.95 seconds between consecutive association pairs. During retrieval, participants were shown the first image of an association pair and instructed to move a cursor to indicate the location of the second image and confirm it. All image pairs were tested in randomized order during each retrieval phase (**Fig. 7**). At regular intervals of 60-90 seconds, the encoding and retrieval tasks were briefly interrupted by a baseline task, in which participants were instructed to fixate a moving white cross on a black background. To assess memory retention, a final retrieval phase was administered after the second post-learning MRI session (t2). The task was implemented in MATLAB R2015b using the Cogent 2000 toolbox.

**Fig. 7.**
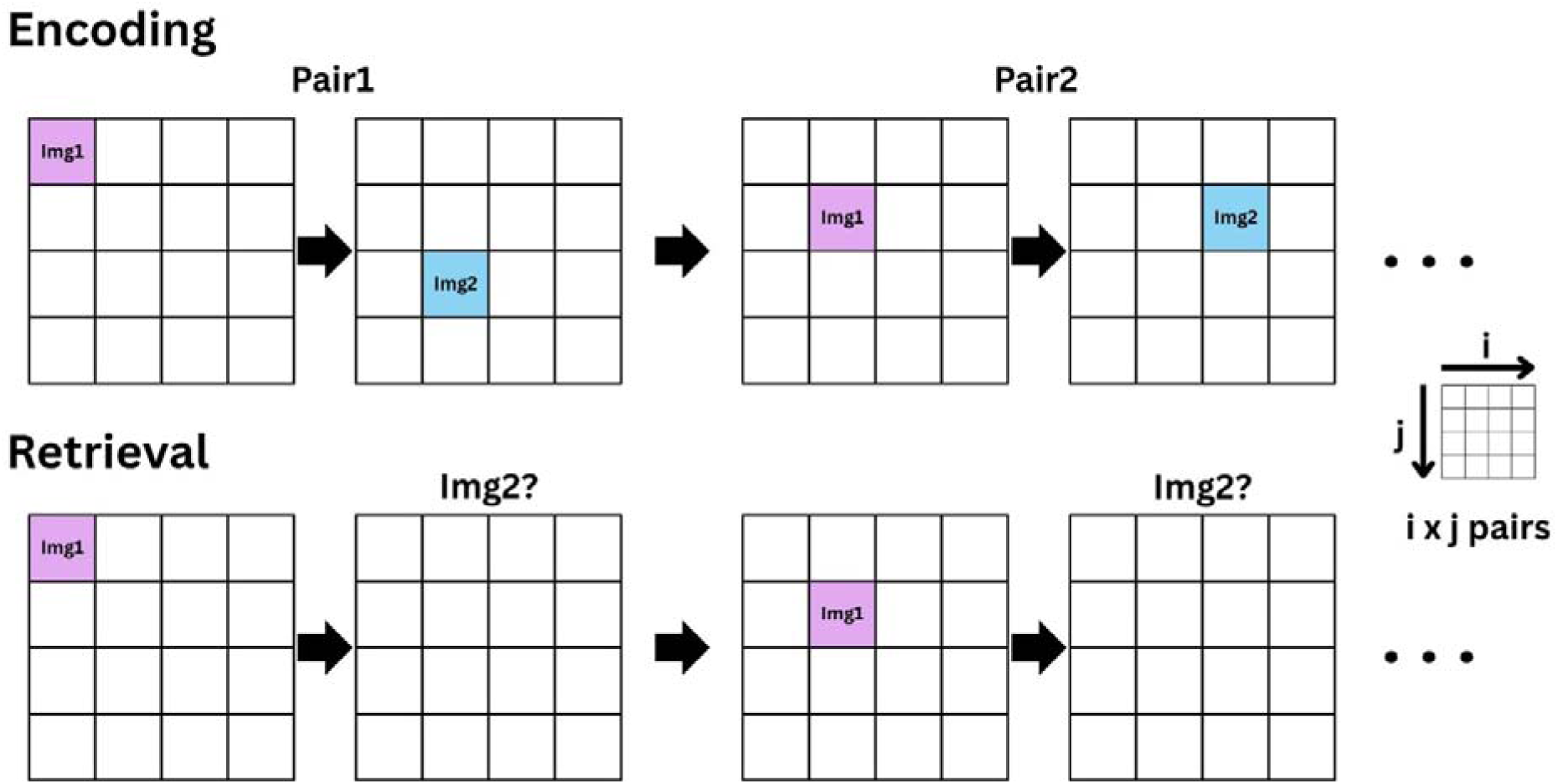
Image–location association task. Schematic illustration of the encoding and retrieval phases of the image-location task. During the encoding phase, participants learned associations between pairs of images and their spatial locations within a grid (either an 8 x 5 grid; Study 1, or a 7×4 grid; Studies 2 and 3). Image pairs were presented sequentially, with each image appearing in a unique grid position. Each image was shown twice per encoding phase, once as the first image (Img1) and once as the second image (Img2). During the retrieval phase, memory for the learned associations was tested by presenting the first image of each pair (Img1) in its original learned location and requiring participants to recall the spatial location of its associated partner image (Img2).

#### Studies 2 and 3

28 images were randomly placed on a 7-by-4 grid. Each participant saw images from one of the following four classes: animate objects, inanimate objects, outdoor scenes, indoor scenes. Each image was part of two different association pairs as either the first or the second image of the pair, resulting in a total of 28 distinct association pairs. During encoding, participants were presented with a random sequence of all association pairs. Each image was shown full screen for 1.5 seconds, before transitioning to the assigned grid location over 1 s, where it remained visible for 3 seconds. Consecutive pairs were separated by an inter-trial interval of 1.7–2.0 seconds. During retrieval, the first image of each pair was presented as a cue for 2.5 seconds, after which participants selected the spatial location of the corresponding second image (**Fig. 7**). All image pairs were tested in randomized order during each retrieval phase. To assess memory retention, a final retrieval phase was administered after the post-learning MRI session. The task was programmed using PsychoPy version 2022.2.4.

### MRI acquisition

In all studies, subjects were instructed to remain still and not to fall asleep during the acquisition period. High-resolution T1-weighted MPRAGE images were acquired with study-specific parameters. Study 1 was conducted with a 3T Siemens PRISMA MRI scanner (Siemens Medical Systems, Erlangen, Germany) using a standard 20-channel head volume coil at the Max Planck Institute for Biological Cybernetics in Tübingen, Germany. The structural image was recorded using the following parameters: *TR* = 5000 ms, *TE* = 2.98 ms, *TI1/TI2* = 700/2500 ms, flip angles of 4° and 5°, a field of view of 256 mm, an isotropic voxel size of 1 mm, 176 sagittal slices, and a GRAPPA acceleration factor of 3. Similarly, study 2 was measured on a 3 T Siemens PRISMA MRI scanner at the University of Freiburg (Medizinische Klinik, Freiburg, Germany) using a standard 64-channel head volume coil, while neuroimaging data of study 3 was acquired on a 3 T Siemens Cima.X scanner equipped with a 64-channel head volume coil. In studies 2 and 3, identical parameters were used, with *TR* = 2300 ms, *TE* = 4.18 ms, *TI* = 900 ms, a flip angle of 9°, a field of view of 256 mm, an isotropic voxel size of 1 mm, 176 sagittal slices, and a GRAPPA acceleration factor of 2. To ensure consistent positioning of measurement volumes across baseline and post-learning sessions and to minimize offline warping, an auto-align scout was used in all studies.

### MRI preprocessing

#### Segmentation and registration

After individual visual inspection, all T1W images were preprocessed using the longitudinal VBM pipeline implemented in the computational anatomy toolbox (CAT12.9; ver. 2550; Gaser et al., 2024) within the statistical parametric mapping software (SPM12; ver. 7771; Friston, 2003). It includes intra-subject realignment, bias correction, segmentation into three tissue types (gray matter, white matter, and cerebrospinal fluid) and non-linear spatial registration to MNI space using DARTEL (Ashburner, 2007). By default, the images are resampled to a voxel size of 1.5 mm (isotropic) during preprocessing. After preprocessing, the quality of each image was assessed through the noise-contrast-ratio (NCR), a quality parameter computed by the CAT12 toolbox. The NCR is calculated based on the standard deviation of the optimized white matter segmentation and the minimum tissue contrast, where higher NCR values indicate lower quality. NCR was extracted for each image and z-transformed across participants within each study. Participants with preprocessing quality scores (z-transformed NCR) greater than 3 were excluded from further analysis (**Fig. 6**).

### ComBat-based feature harmonization

We used the neuroCombat harmonization procedure to reduce possible effects of site (Johnson et al., 2007). The harmonization model expects the imaging features to be a linear combination of site effects and biological variables. Here, ComBat harmonization was implemented voxel-wise across the entire gray matter mask via a custom Python script using the default settings (eb = True, parametric = True, mean_only = False, ref_Batch = None) and defining the condition (learning, control) and the time since the baseline scan (0 hours, 2 hours, 12 hours) as covariates. This procedure was applied prior to statistical analysis and was designed to reduce scanner-related variability while preserving effects of condition and time. After harmonization, the images were smoothed with a Gaussian kernel at 8 mm full-width at half maximum (FWHM). The segmented, modulated, harmonized, and smoothed GM images were then used for further statistical analysis.

### Statistics and Reproducibility

#### Behavioral performance

Performance on the image-location task was assessed as the percentage of correctly recalled locations in each retrieval phase (Brodt et al., 2018). Repetitions R1–R4 occurred during the initial encoding-retrieval task block within the same session. The fifth retrieval (R5) was a separate, final retrieval assessed after the post-learning MRI session (∼2 h post-learning in Study 2; ∼12 h post-learning in Studies 1 and 3). We extracted performance from all four within-block repetitions (R1–R4), as well as from this fifth retrieval (R5). Memory retention was defined as the difference between R4 and R5 (R5–R4), capturing the change in performance across the post-learning consolidation interval relative to performance at the end of the initial encoding–retrieval block. A two-way mixed ANOVA with factors of repetition and study site was used to assess learning curves and potential between-study differences in performance. In the post-hoc analysis, significant main effects were addressed using post hoc t-tests (two-tailed) with Bonferroni correction. Because the interval between R4 and R5 differed across studies (∼2 h in Study 2 versus ∼12 h in Studies 1 and 3), we additionally compared R4 and R5 accuracy separately within each study using paired t-tests.

#### VBM group-level analysis

Group-level GMV changes were modeled using a flexible factorial design with the factors of *subject*, *time* and *group* to assess main effects of time and group as well as their interaction. All images were organized in a factor matrix with four columns: (1) a constant column of ones as required by CAT12, (2) a unique subject identifier (“1” to “198”), (3) a factor representing condition (learning or control), (4) a factor representing time (baseline, 2 h, or 12 h). A thresholded masking at an absolute value of 0.1 was applied in the model, without any additional masking. As a result, the design matrix included: time1 × condition1, time1 × condition2, time2 × condition1, time2 × condition2, time3 × condition1, time3 × condition2, subject1, …subject198. We proceeded with t-contrasts to assess learning-induced volumetric changes: learning > control from baseline to 2 h (1 -1 -1 1 0 0), learning > control from baseline to 12 h (1 -1 0 0 -1 1), and control > learning from 2 h to 12 h (0 0 -1 1 1 -1). Significant results are reported at an uncorrected threshold of *p* < 0.001 with a minimum cluster extent of *k* = 20 voxels given the small and transient effect sizes expected in learning-induced GMV changes (see Villa et al., 2026). FWE-corrected results at the peak level are reported additionally where surviving clusters are present. Lastly, we used the conjunction analysis implemented in SPM12 (Friston, 2003) to identify the overlap between the clusters obtained from the t1-t0 and the t2-t1 interaction contrasts described above. The extent of spatial overlap was further quantified by the Dice coefficient (Dice, 1945; Moore et al., 2023; Shen et al., 2013), which ranges from 0 (no spatial overlap) to 1 (complete overlap) and was computed as 2 × (overlap voxels) / (n(t0t1) + n(t1t2)). The dice coefficient should be interpreted alongside the reported overlap voxel counts and cluster sizes, as only clusters of identical size can have a dice coefficient of 1. For all clusters that overlapped between the t1-t0 and the t2-t1 contrast, cluster labels were extracted from the probabilistic HO Cortical and Subcortical Atlases, as distributed with FSL, both for the peak coordinate and, additionally, based on the mean likelihood across all voxels within each cluster (labels > 1% were reported). The same procedure was applied using the deterministic AAL atlas distributed with CAT12, with the proportion of voxels assigned to each label reported instead of probabilistic likelihoods.

#### Cluster-level GMV change over time

To quantify GMV change within clusters identified by the VBM analysis, mean voxel values were extracted from the GMV maps of all subjects. Because these clusters were defined based on the whole-brain VBM analysis, the following region-level analysis should be interpreted as non-independent characterization of the identified clusters rather than as independent confirmation of the VBM findings. GMV change was operationalized as the difference in mean cluster value between two timepoints (later minus earlier). Given the nested structure of the data combining within-subject (studies 1 and 3) and between-subject (study 2) designs, a linear mixed effect model (LMM) was fitted separately for each cluster of interest to assess short-term GMV changes (t1-t0). Based on the recommendation (Forstmeier & Schielzeth 2011; Uhlig et al., 2023), we compared a null model and a full model that were identical except for the term of interest. In both models, GMV change served as the dependent variable, with study sites (studies 1 and 2) and condition (learning, control) as fixed factors and subject as a random factor with random intercepts only. The null model included main effect of study and condition (*GMV change ∼ study site + condition + (1/subject)*), while the full model included the interaction between conditions (*GMV change ∼ study site x condition + (1/subject)*). Both models were fitted via maximum likelihood using the MixedLM function in Python, and model fit was compared using a χ² test of log-likelihood values, with significance set at α = .05. The pairwise post-hoc comparisons between conditions were performed using estimated marginal means using R (ver. 4.5.3, toolboxes: *emmeans* 2.0.2, lme4 2.0.1). As only study 1 contributed data for the t2-t1 interval (*n*=32), given its within-subject design, a two-tailed paired t-test was used for assessing GMV changes. The contributing studies and sample sizes for this and subsequent analyses are listed in **Table S3**.

#### Association between memory retention and GMV

Memory retention was defined as the difference in performance accuracy between fourth and fifth retrieval. Pearson correlations were computed to examine associations between GMV change in bilateral PC at both the t1-t0 and t2-t1 intervals and memory retention. Due to differences in study design, only study 2 contributed to the correlation analysis for the t1-t0 interval, as it included the final retrieval required to compute memory retention. In contrast, only study 1 provided MRI measurements at both t1 and t2, along with a final retrieval following t2, and was therefore used for the t2-t1 analysis. To further characterize this relationship, participants were additionally divided into two groups using GMV by median split and memory retention was compared between groups using an independent-samples t-test.

#### The effect of sleep on learning-related GMV changes

The present dataset included both sleep and wake intervals between scanning sessions, allowing us to examine whether sleep modulated learning-induced GMV changes. Study 1 included two subgroups: a sleep group, scanned in the evening and sleeping between t1 and t2, and a wake group, scanned in the morning and remaining awake between these timepoints. Because only Study 1 provided both groups, the analysis was limited to this dataset. For each parietal cortex cluster, we performed a two-way mixed ANOVA with the factors sleep (sleep vs. wake) and condition (learning vs. control). Follow-up comparisons between learning and control within each subgroup were conducted using two-tailed pairwise t-tests with Bonferroni correction for multiple testing. To examine sleep effects across the full post-learning interval (t2-t0), the same analytical approach was applied. Given that no significant whole-brain clusters emerged for this contrast, GMV change was extracted from clusters defined by the spatial conjunction of the early learning increase (t1-t0) and subsequent renormalization (t1–t2) contrasts, using the conjunction procedure in SPM12. For this interval, both studies 1 and 3 contributed data to the sleep group and employed within-subject designs; therefore, a two-way mixed ANOVA was applied directly without requiring an LMM. A publication on sleep–wake effects on behavioral performance in study 1 is currently available as a preprint (Brodt et al., 2023).

All statistical analyses were performed using Python (ver. 3.12, toolboxes: pingouin 0.6.0, statsmodels 0.14.6, Scipy 1.17.0) and R (ver. 4.5.3., toolboxes: emmeans 2.0.2, lme4 2.0.1).

## Data availability

The preprocessed T1W-MRI images and image-localization task results for all participants have been deposited on the open science framework platform (OSF): osf.io/pu78e. All (unthresholded) statistical maps are available at (Gorgolewski et al., 2015) for detailed inspection in 3D (https://neurovault.org/collections/AOJBHIWN).

## Code Availability

The code employed for the analyses of the data is available at the author’s open science framework platform (OSF): osf.io/pu78e. The software and tools used include: MATLAB R2023b (Mathworks), python (ver. 3.12).

## Author contributions

**Felix Kleinschroth:** Conceptualization, Methodology, Data collection (study 3), Formal analysis, Data curation, Writing – original draft, Writing – review & editing, Visualization. **Stella Villa:** Data collection (study 3), Writing – review & editing. **Antonia Lenders:** Data collection (study 2), Writing – review & editing. **Svenja Brodt:** Data collection (study 1), Writing – review & editing. **Steffen Gais:** Data collection (study 1), Writing – review & editing. **Monika Schönauer:** Data collection (study 1), Writing – review & editing. **Deniz Kumral:** Conceptualization, Investigation, Data Collection (studies 2,3), Writing – original draft, Writing – review & editing, Resources, Supervision.

## Acknowledgements

The authors want to thank all participants and students supporting the data collection, as well as the CoreFacility ‘Magnetic Resonance Development and Application Center Freiburg (MRDAC)’, Department of Radiology University Medical Center Freiburg, Faculty of Medicine, University of Freiburg, for support in acquisition of the data. We further thank Arno Villringer from the Max-Planck-Institute Leipzig for fruitful discussion on the biology underlying observed MR signal changes.

## Supplementary Material

**Tab. S1.** Interaction – GMV increase (t0 vs t1) x (learning vs control)

| Region | *n* voxels | $T_{\text{peak}}$ | $x_{\text{peak}}$ | $y_{\text{peak}}$ | $z_{\text{peak}}$ |
| --- | --- | --- | --- | --- | --- |
| Right Parietal Operculum Cortex / Planum Temporale* | 123 | 4,77 | 40 | -36 | 21 |
| Left Parietal Operculum Cortex / Planum Temporale | 134 | 4,49 | -44 | -42 | 20 |
| Left Cerebellum | 270 | 4,31 | -33 | -74 | -26 |
| Left Lateral Occipital Cortex | 128 | 3,57 | -27 | -81 | 32 |
| Left Anterior Medial Temporal Lobe | 69 | 3,72 | -27 | 3 | -15 |
| Left Precentral Gyrus | 20 | 3,50 | -12 | -21 | 51 |

Regions listed exhibited a significant peak voxel effect at $p_{\text{uncorr}} < .001$ , $df = [1.0, 444.0]$ and had a cluster size of $k \geq 20$ . \* Regions also survived peak-level FWE correction in the whole brain analysis at $p_{\text{FWE}} < .05$ . Peak voxel MNI coordinates X,Y and Z are given in mm.

**Tab. S2.** Interaction – GMV decrease (t1 vs t2) x (learning vs control)

| Region | n voxels | T<sub>peak</sub> | X<sub>peak</sub> | Y<sub>peak</sub> | Z<sub>peak</sub> |
| --- | --- | --- | --- | --- | --- |
| Right Parietal Operculum<br>Cortex / Planum Temporale* | 195 | 5,02 | 39 | -34 | 18 |
| Left Parietal Operculum<br>Cortex / Planum Temporale | 164 | 4,45 | -39 | -39 | 18 |
| Left Anterior Medial<br>Temporal Lobe | 250 | 4,42 | -22 | 3 | -15 |
| Left Subcallosal Cortex | 33 | 3,64 | -6 | 28 | -10 |
| Right Lateral Occipital Cortex | 30 | 3,81 | 40 | -63 | 14 |
| Left Cerebellum | 144 | 3,66 | -9 | -84 | -27 |
| Left Supramarginal Gyrus | 30 | 3,53 | -51 | -26 | 26 |
| Right Temporal Pole | 26 | 3,66 | 42 | 12 | -27 |
| Left Precentral Gyrus | 22 | 3,38 | -12 | -21 | 51 |
| Left Cerebellum | 57 | 3,45 | -24 | -80 | -24 |
| Left Cerebellum | 31 | 3,28 | -16 | -70 | -18 |
| Right Cingulate Gyrus | 65 | 3,29 | 3 | 32 | 15 |

Regions listed exhibited a significant peak voxel effect at $p_{\text{uncorr}} < .001$ , $df = [1.0, 444.0]$ and had a cluster size of $k \geq 20$ . \* Regions also survived peak-level FWE correction in the whole brain analysis at $p_{\text{FWE}} < .05$ . Peak voxel MNI coordinates X,Y and Z are given in mm.

**Tab. S3.** Sample size and study contributions for each contrast.

| Time points | Time interval (h) | Contributing Studies | N | Group size (learning / control) | Establishes |
| --- | --- | --- | --- | --- | --- |
| t1-t0 | 0 - 2 | 1, 2 | 149 | 109 / 62 | Early expansion |
| t2-t1 | 2 - 12 | 1 | 37 | 37 / 32 | renormalization |
| t2-t0 | 0 - 12 | 1, 3 | 96 | 96 / 91 | net trajectory |
| t2-t1 (sleep) | 2 - 12 | 1 | 32 | 17 / 15 (sleep / wake) | sleep modulation of renormalization |

*Note.* The sample size and contributing studies for each timepoint combination reported in the manuscript differ due to differences in study design. This table lists the sample size, contributing studies, and group sizes for each contrast, along with the interpretive role each analysis plays in the Discussion. The 32 subjects contributing to the sleep analysis are the same individuals from the t2-t1 control group. See methods section for exclusion details.

**Tab. S4.** Conjunction between the interaction clusters.

| Area | $n$ (t0t1) | $n$ (t1t2) | $n$ (overlap) | dice coefficient |
| --- | --- | --- | --- | --- |
| Right Parietal Operculum Cortex | 123 | 195 | 91 | 0.572 |
| Left Parietal Operculum Cortex | 134 | 164 | 119 | 0.799 |
| Left Anterior Medial Temporal Lobe | 69 | 250 | 25 | 0.157 |

*Note.* Regions listed had overlapping clusters of size $k \geq 20$ between the interactions from **Tab. S1** and **Tab. S2**.

**Tab. S5.**
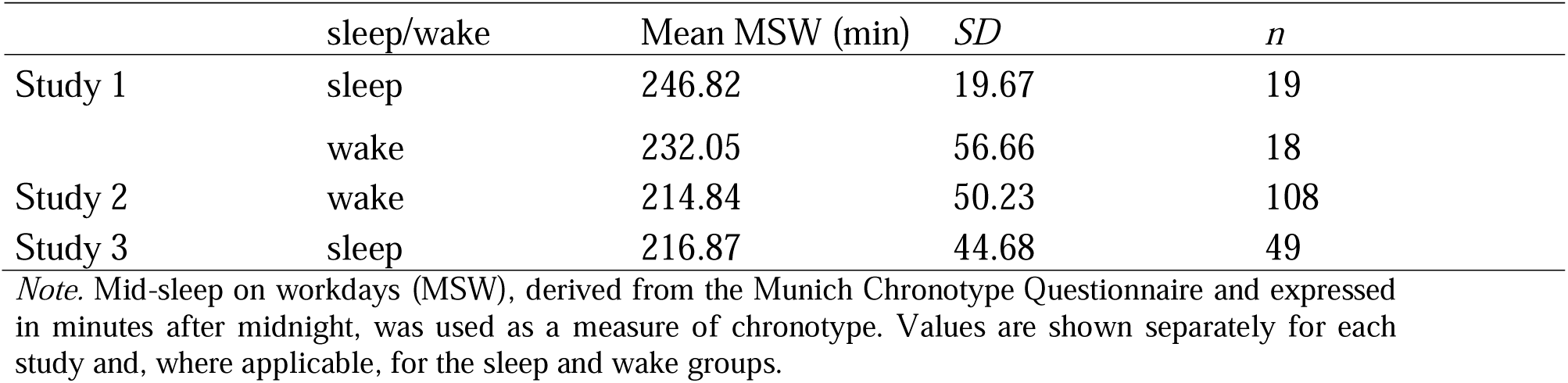
Mean values of the Munich Chronotype Questionnaire for each study.

|  | sleep/wake | Mean MSW (min) | *SD* | *n* |
| --- | --- | --- | --- | --- |
| Study 1 | sleep | 246.82 | 19.67 | 19 |
|  | wake | 232.05 | 56.66 | 18 |
| Study 2 | wake | 214.84 | 50.23 | 108 |
| Study 3 | sleep | 216.87 | 44.68 | 49 |

*Note.* Mid-sleep on workdays (MSW), derived from the Munich Chronotype Questionnaire and expressed in minutes after midnight, was used as a measure of chronotype. Values are shown separately for each study and, where applicable, for the sleep and wake groups.

**Tab. S6.** Labels for PC clusters.

| Atlas | Type | Contrast | Cluster | Labels (Cluster) | Labels (Peak) |
| --- | --- | --- | --- | --- | --- |
|  |  |  |  |  | t1-t0, left: [-44, -42, 20]<br>t2-t1, left: [-39, -39, 18]<br>t1-t0, right: [40, -36, 21]<br>t2-t1, right: [39, -34, 18] |
| Harvard-Oxford<br>Cortical Atlas | Probabilistic | t1-t0 | Left PC | Parietal Opercular Cortex (22.6%),<br>Planum Temporale (13.3%),<br>Supramarginal Gyrus (6.9%),<br>Superior Temporal Gyrus (1.9%) | Planum Temporale (16.0%),<br>Parietal Opercular Cortex (11.0%),<br>Supramarginal Gyrus (8.0%),<br>Superior Temporal Gyrus (4.0%) |
|  |  |  | Right PC | Parietal Opercular Cortex (34.7%),<br>Planum Temporale (10.1%),<br>Supramarginal Gyrus (4.6%) | Parietal Opercular Cortex (28.0%),<br>Supramarginal Gyrus (12.0%),<br>Planum Temporale (2.0%) |
|  |  | t2-t1 | Left PC | Parietal Opercular Cortex (26.6%),<br>Planum Temporale (17.9%),<br>Supramarginal Gyrus (5.9%),<br>Superior Temporal Gyrus (1.9%) | Parietal Opercular Cortex (32.0%),<br>Planum Temporale (20.0%),<br>Supramarginal Gyrus (7.0%),<br>Superior Temporal Gyrus (3.0%) |
|  |  |  | Right PC | Parietal Opercular Cortex (28.3%),<br>Planum Temporale (23.4%),<br>Supramarginal Gyrus (3.8%),<br>Heschl's Gyrus (4.4%) | Parietal Opercular Cortex (30.0%),<br>Planum Temporale (27.0%),<br>Supramarginal Gyrus (5.0%) |
| Harvard-Oxford<br>Subcortical<br>Atlas | Probabilistic | t1-t0 | Left PC | Left Cerebral White Matter (52.7%),<br>Left Cerebral Cortex (46.8%) | Left Cerebral White Matter<br>(55.6%),<br>Left Cerebral Cortex (42.3%) |
|  |  |  | Right PC | Right Cerebral Cortex (50.7%),<br>Right Cerebral White Matter (49.2%) | Left Cerebral Cortex (57.3%),<br>Left Cerebral White Matter (42.7%) |
|  |  | t2-t1 | Left PC | Left Cerebral Cortex (54.1%),<br>Left Cerebral White Matter (44.4%) | Left Cerebral Cortex (62.3%),<br>Left Cerebral White Matter (37.7%) |
|  |  |  | Right PC | Right Cerebral Cortex (62.7%),<br>Right Cerebral White Matter (37%) | Left Cerebral Cortex (63.0%),<br>Left Cerebral White Matter (36.8%) |
| Automated<br>Anatomical<br>Labeling Atlas | Deterministic | t1-t0 | Left PC | Temporal_Sup_L: 76/134 voxels (56.7%),<br>SupraMarginal_L: 23/134 voxels (17.2%),<br>Rolandic_Oper_L: 11/134 voxels (8.2%),<br>Insula_L: 1/134 voxels (0.7%) | -- |
|  |  |  | Right PC | Rolandic_Oper_R: 21/123 voxels (17.1%),<br>Insula_R: 18/123 voxels (14.6%),<br>SupraMarginal_R: 10/123 voxels (8.1%),<br>Temporal_Sup_R: 5/123 voxels (4.1%),<br>Heschl_R: 3/123 voxels (2.4%) | -- |
|  |  | t2-t1 | Left PC | Temporal_Sup_L: 97/164 voxels (59.1%),<br>Rolandic_Oper_L: 24/164 voxels (14.6%),<br>SupraMarginal_L: 15/164 voxels (9.1%),<br>Insula_L: 2/164 voxels (1.2%) | Temporal_Sup_L |
|  |  |  | Right PC | Heschl_R: 47/195 voxels (24.1%),<br>Rolandic_Oper_R: 35/195 voxels (17.9%),<br>Temporal_Sup_R: 26/195 voxels (13.3%),<br>Insula_R: 23/195 voxels (11.8%),<br>Supramarginal_R: 5/195 voxels (2.6%) | -- |

*Note.* Labels for the left and right parietal operculum (PC) clusters based on the VBM analysis for the t1-t0 contrast and the t2-t1 contrast. Labels were derived from the Harvard-Oxford Cortical atlas and Subcortical atlas as well as the Automated Anatomical Labeling atlas on both, a cluster-level and the peak-level.

**Tab. S7.** Labels for Anterior Medial Temporal cluster.

| Atlas | Type | Contrast | Labels (Cluster) | Labels (Peak) |
| --- | --- | --- | --- | --- |
|  |  |  |  | t1-t0: [-27, 3, -15]<br>t2-t1: [-22, 3, -15] |
| Harvard-Oxford Cortical Atlas | Probabilistic | t1-t0 | Frontal Orbital Cortex (5.4%),<br>Parahippocampal Insular Cortex (4.4%),<br>Gyrus (2.0%),<br>Temporal Pole (2.1%) | Parahippocampal Gyrus, anterior division (2.0%) |
|  |  | t2-t1 | Parahippocampal Gyrus (6.4%),<br>Temporal Pole (2.9%),<br>Frontal Orbital Cortex (2.8%) | Parahippocampal Gyrus, anterior division (2.0%) |
| Harvard-Oxford Subcortical Atlas | Probabilistic | t1-t0 | Cerebral Cortex (72.4%),<br>Amygdala (13.3%),<br>Cerebral White Matter (6.3%) | Cerebral Cortex (76.8%),<br>Amygdala (13.6%),<br>Cerebral White Matter (2.5%) |
|  |  | t2-t1 | Cerebral Cortex (39.9%),<br>Amygdala (31.4%),<br>Hippocampus (11.8%),<br>Cerebral White Matter (2.4%) | Cerebral Cortex (78.5%),<br>Amygdala (3.5%) |
| Automated Anatomical Labeling Atlas | Deterministic | t1-t0 | Olfactory_L: 3/69 voxels (4.3%),<br>Insula_L: 2/69 voxels (2.9%),<br>Amygdala_L: 8/69 voxels (11.6%) | Amygdala_L |
|  |  | t2-t1 | Amygdala_L: 96/250 voxels (38.4%),<br>Olfactory_L: 46/250 voxels (18.4%),<br>Hippocampus_L: 45/250 voxels (18.0%),<br>Temporal_Pole_Sup_L: 27/250 voxels (10.8%),<br>ParaHippocampal_L: 2/250 voxels (0.8%) | Left Olfactory Cortex |

*Note.* Labels for the left anterior medial temporal cluster based on the VBM analysis for the t1-t0 contrast and the t2-t1 contrast. Labels were derived from the Harvard-Oxford Cortical atlas and Subcortical atlas as well as the Automated Anatomical Labeling atlas on both, a cluster-level and the peak-level.

**Tab. S8.** Labels for Anterior Medial Temporal cluster conjunction (t1-t0 ∩ t2-t1)

| Atlas | Type | Labels (Cluster) | Labels (Peak – [-26, 3, -15]) |
| --- | --- | --- | --- |
| Harvard-Oxford Cortical Atlas | Probabilistic | Frontal Orbital Cortex (3.8%),<br>Parahippocampal Gyrus (2.4),<br>Temporal Pole (1.5%) | Parahippocampal Gyrus,<br>anterior division (3.0%) |
| Harvard-Oxford Subcortical Atlas | Probabilistic | Cerebral Cortex (73.4%),<br>Amygdala (12.8%),<br>Cerebral White Matter (3.0%) | Cerebral Cortex (78.6%),<br>Amygdala (12.8%),<br>Cerebral White Matter (1.8%) |
| Automated Anatomical Labeling Atlas | Deterministic | Amygdala_L: 3/25 voxels (12.0%),<br>Olfactory_L: 3/25 voxels (12.0%) | Amygdala_L |

*Note.* Labels for the cluster resulting from the conjunction analysis between the left anterior medial temporal cluster from the the t1-t0 contrast and the t2-t1 contrast. Labels were derived from the Harvard-Oxford Cortical atlas and Subcortical atlas as well as the Automated Anatomical Labeling atlas on both, a cluster-level and the peak-level.

**Fig. S1.**
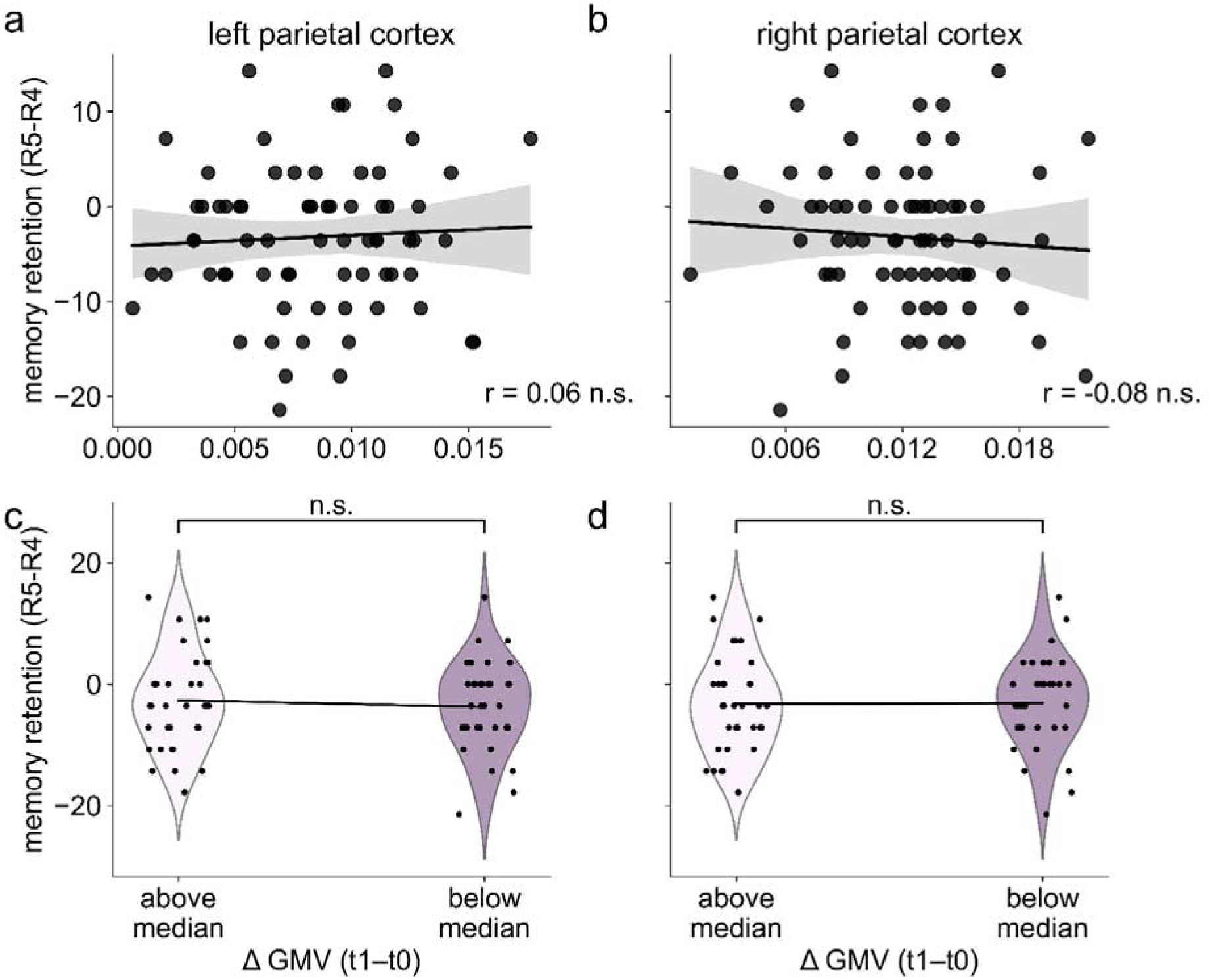
Association between memory retention and short-term GMV changes (t1-t0) in the parietal cortex. We assessed the association between GMV changes (t1-t0) and memory retention in the left (**a**) and right (**b**) parietal cortex (PC) clusters using Pearson correlation and observed no significant correlation in the left PC (*r* = .055, 95% *CI* = [-0.18, 0.29], *p* = .65, *n* = 71) or right PC (*r* = -.076, 95% *CI* = [-0.30, 0.16], *p* = .76, *n* = 71). When comparing memory retention after splitting the data by median GMV change (t1 – t0) in the left **(c,** above: *M* = -2.68, *SD* = 7.89, *n* = 36; below: *M* = -3.78, *SD* = 7.55, *n* = 35) and right PC **(d,** above: *M* = -3.27, *SD* = 7.93, *n* = 36; below: *M* = -3.16, *SD* = 7.54, *n* = 35)), we observed similarly no significant differences for either the left (*t*(69) = -0.60, *d* = -0.14, *p* = .55) or right PC (*t*(69) = 0.06, *d* = -0.014, *p* = .95). Individual data points are shown over the violin plots. GMV, gray matter volume; R4, fourth retrieval; R5, fifth retrieval; PC, parietal cortex; t0, baseline; t1, 2 h post-learning. Gray, shaded regions indicate the 95 % confidence interval; Bonferroni-corrected; n.s. not significant.

**Fig. S2.**
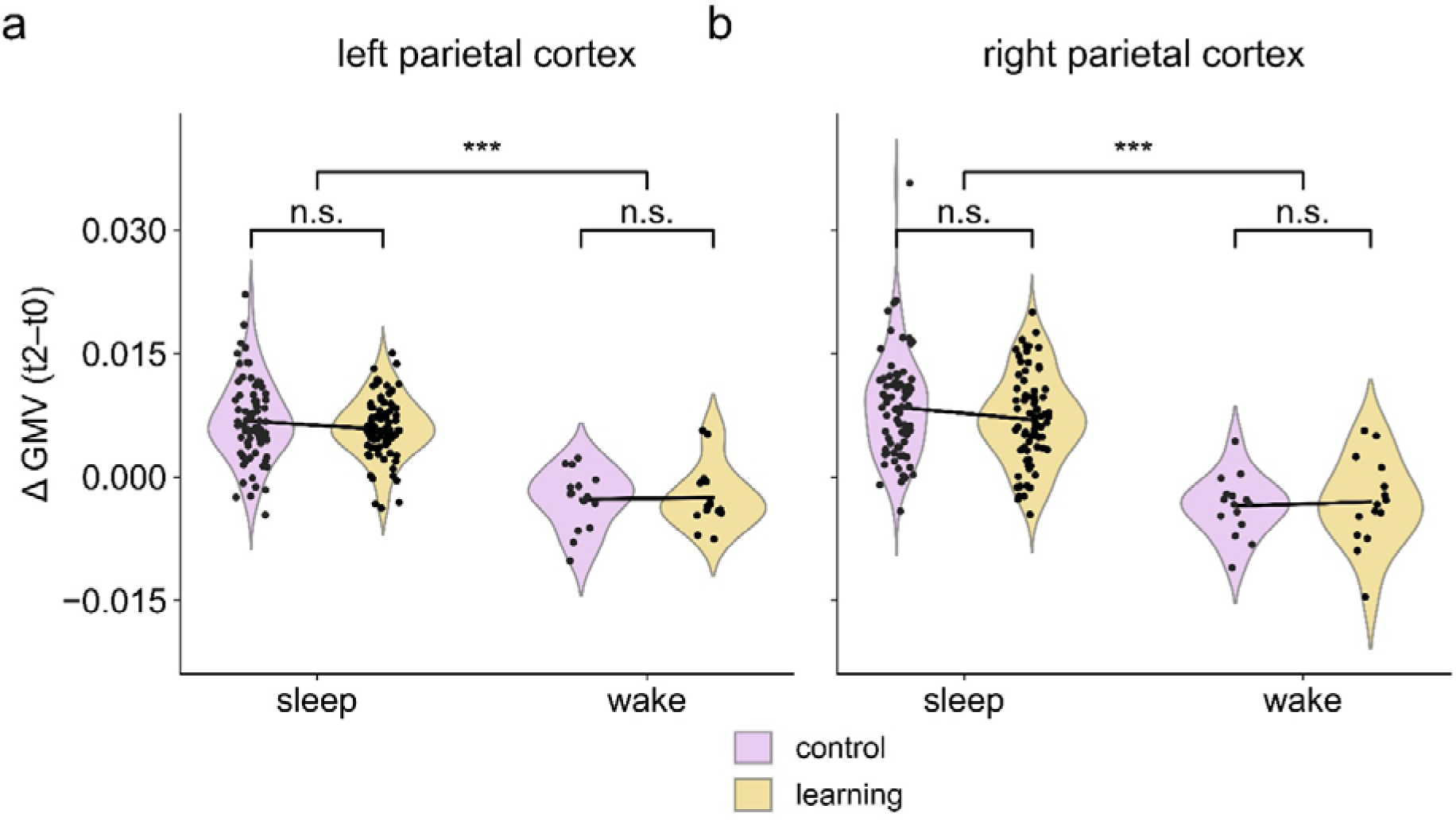
Effect of sleep on parietal long-term GMV change (t2-t0). GMV change between baseline and 12 h post-learning (t2-t0) in the left (**A.**) and right (**B.**) parietal operculum cortex (PC), stratified by sleep condition (sleep group: studies 1 and 3, *n* = 76; wake group: study 1 only, *n* = 15). The sleep group showed significantly greater GMV retention compared to the no-sleep group in both the left PC (**a**; sleep: *M* = 0.006, *SD* = 0.004; wake: *M* = −0.003, *SD* = 0.004) and right PC (**b**; sleep: *M* = 0.007 *SD* = 0.005; wake: *M* = −0.003, *SD* = 0.004). A 2 × 2 mixed ANOVA confirmed a significant main effect of sleep left: *F*(1, 89) = 82.27, *p* < .001, η² = 0.48; right: *F*(1, 89) = 76.34, *p* < .001, η² = 0.46); neither the main effect of condition nor the sleep × condition interaction were significant. A post-hoc comparison of differences between conditions was not found significant in either PC cluster. GMV, gray matter volume; t0, baseline pre-learning; t2, 12 h post-learning; Bonferroni-corrected; *** p < .001, n.s. not significant.

**Fig. S3.**
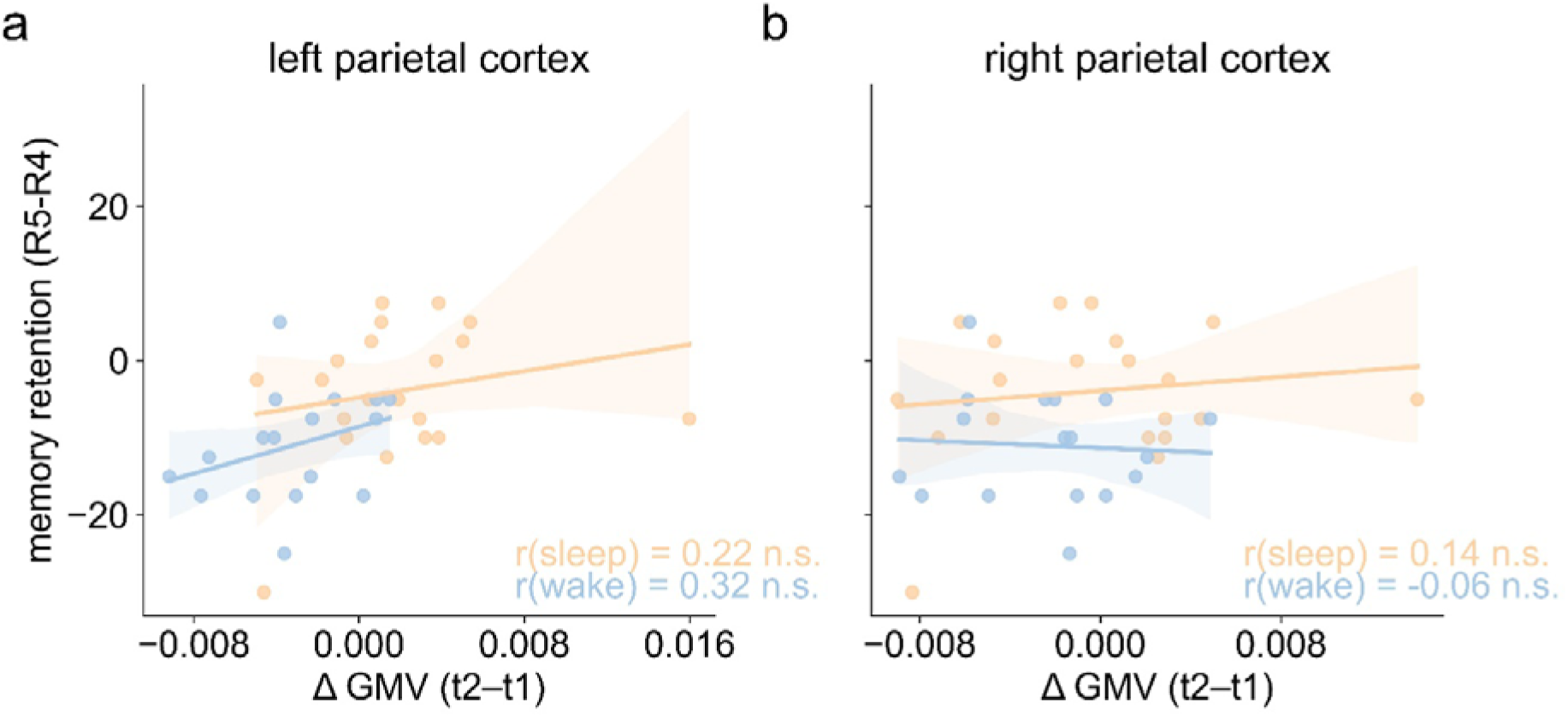
Relationship between parietal GMV renormalization and memory retention by sleep and wake groups. We computed the Pearson correlation separately for wake (blue) and sleep (orange) groups and observed no association between memory retention and GMV change (t2-t1) in the left parietal operculum cortex (PC; **a**; sleep: *r* = .22, 95% *CI* = [-0.25, 0.60], *p* = .16; wake: *r* = .32, 95% *CI* = [-0.19, 0.68], *p* = 0.20) and in the right PC (**b**; sleep: *r* = .14, 95% *CI* = [-0.32, 0.55], *p* = .55; wake: *r* = -.06, 95% *CI* = [-0.59, 0.43], *p* = .81). Shaded areas indicate the 95 % confidence interval. GMV, gray matter volume; R4, fourth retrieval; R5, fifth retrieval; t1, 2 h post-learning; t2, 12 h post-learning; n.s. not significant.

**Fig. S4.**
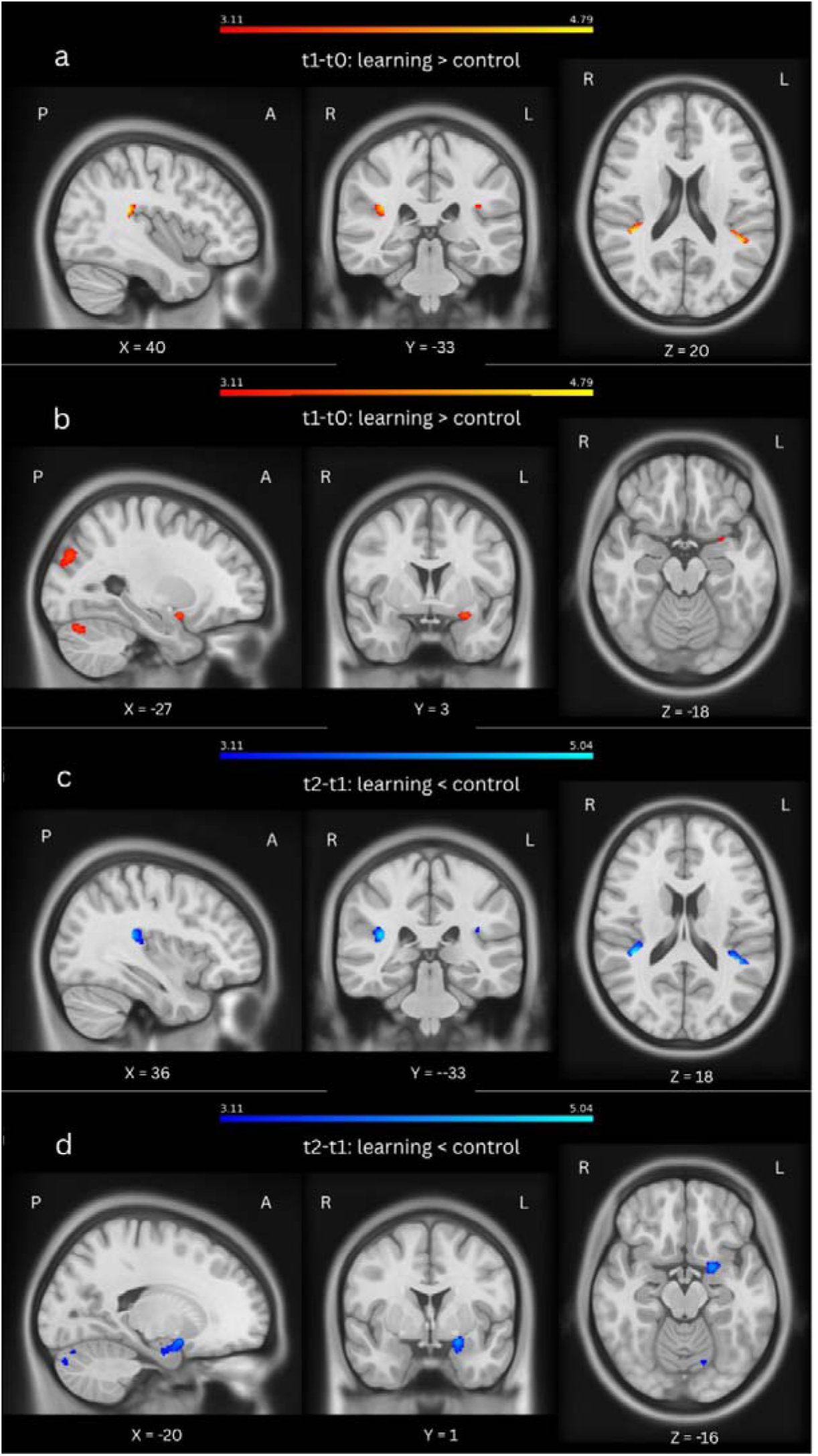
Parietal and anterior Medial Temporal clusters in MNI space. Voxel-based morphometry identified clusters in the parietal operculum (PC) region extending to Planum Temporale and left anterior Medial Temporal Lobe associated with the initial increase in gray matter volume (GMV; t1–t0; **a,b**) and the subsequent decrease (t2–t1; **c,d)** in the learning condition. While first and third rows (a-c) show the PC clusters and second and forth rows (b-d) show left anterior Medial Temporal cluster (see also **Tables S1 and S2**).

**Fig. S5.**
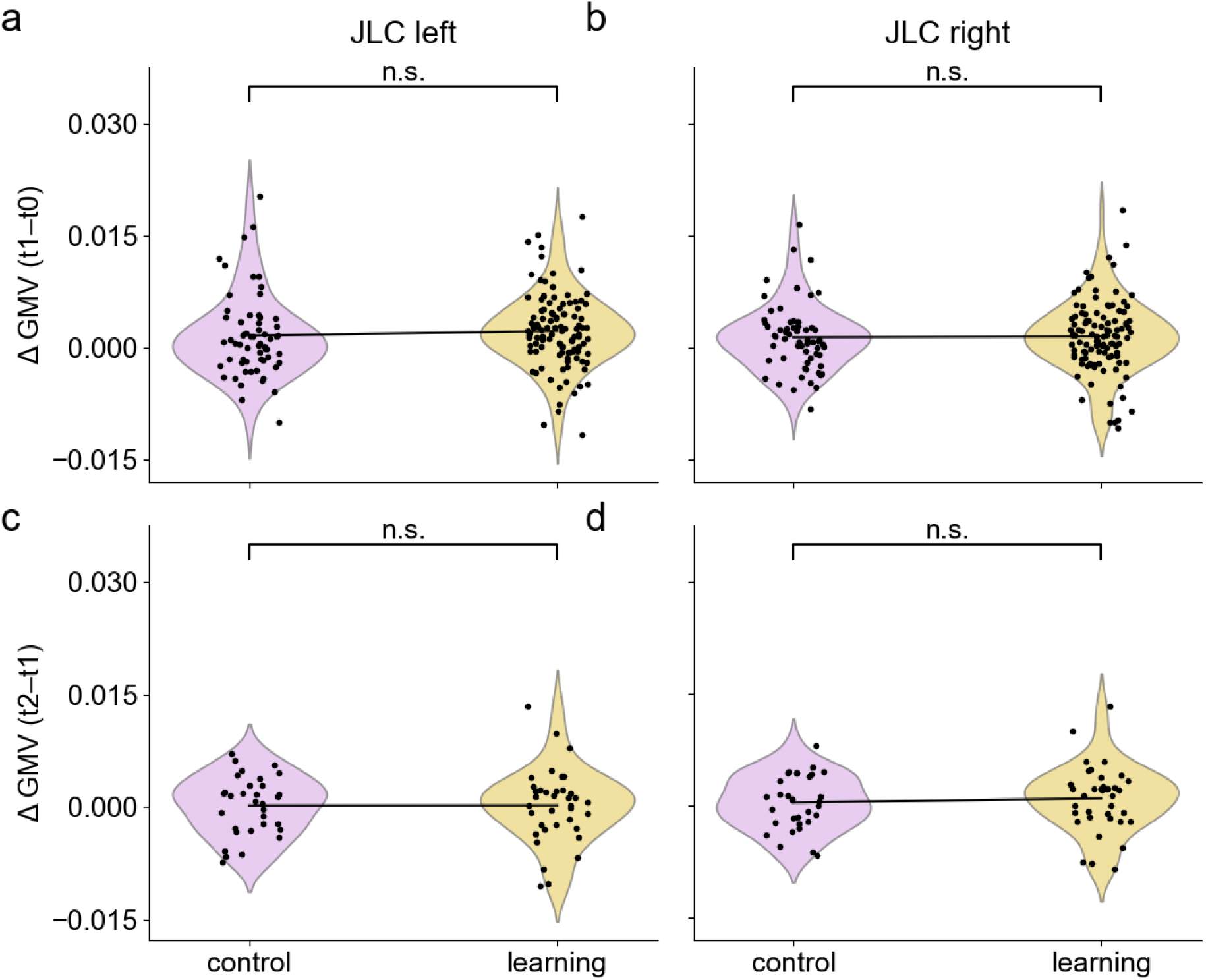
Control analysis analogous to. Fig. 3**. using the juxtapositional lobule cortex (JLC).** GMV change (t1-t0) in the left (p_uncorr_ < 0.001, k > 20 voxels) and right JLC control region for learning (yellow) and control (purple) conditions across studies 1 and 2 (*n*_learning_ = 109, *n*_control_ = 62). (a-b). For the GMV change from baseline to 2 h post-learning (t1-t0), we observed no differences between the control and the learning condition (left: β = -0.00005, *SE* = 0.0008, *p* = .95; right: β = 0.0004, *SE* = 0.0007, *p* = .55). (c-d). Likewise, GMV change from 2 h to 12 h post-learning (t2-t1), paired t-tests (*n*_learning_ = 32, *n*_control_ = 32) indicated no difference between the two groups on either left (*t*(31) = -0.07, *d* = -0.018, *p* = .95) or right side (*t*(31) = -0.74, *d* = -0.19, *p* = .47). Individual data points are plotted on top of the violin plots. GMV, gray matter volume; t0, baseline; t1, 2 h post-learning; t2, 12 h post-learning (Bonferroni-corrected; n.s. not significant)

**Fig. S6.**
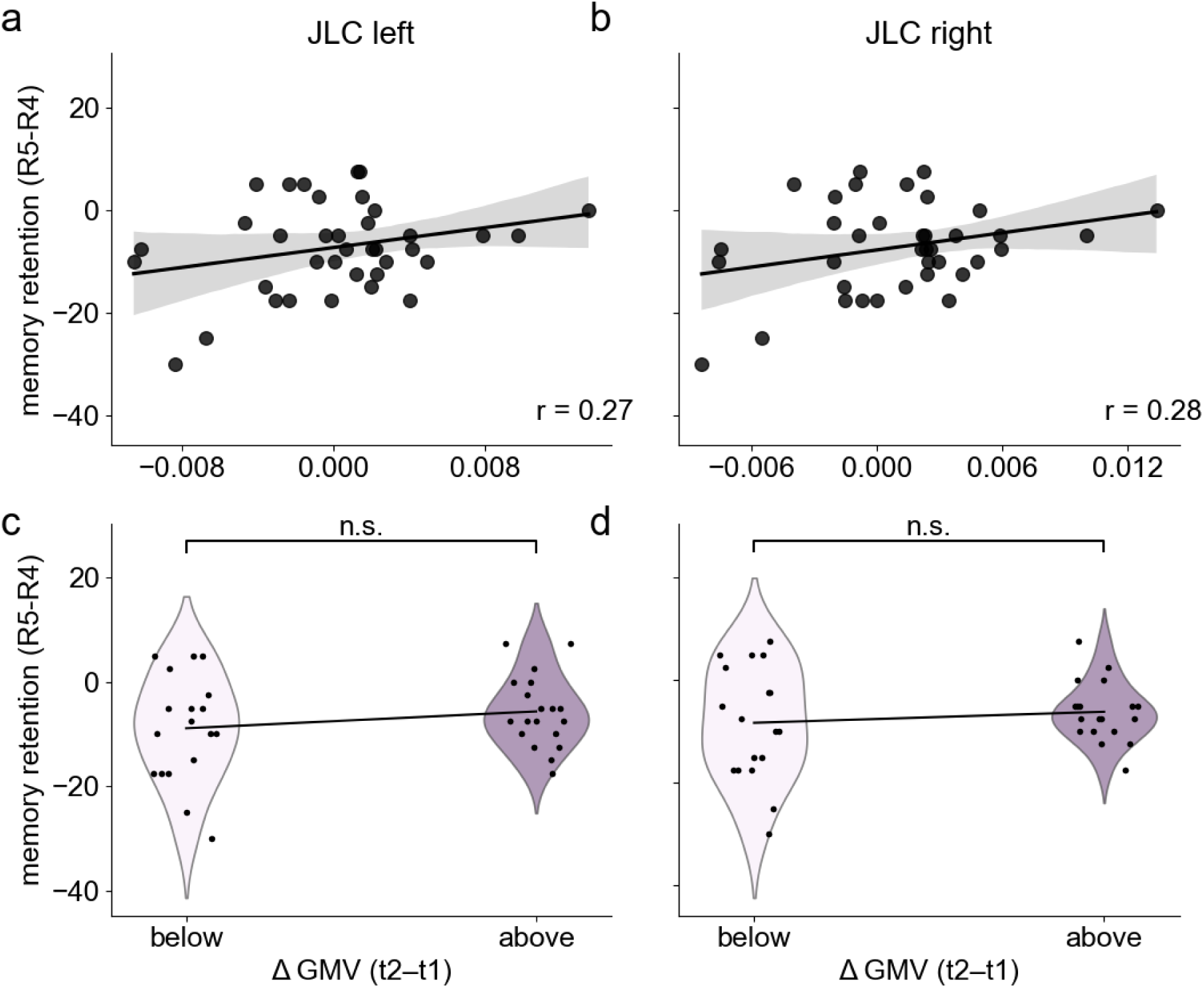
Control analysis analogous to. Fig. 4**. using juxtapositional lobule cortex (JLC).** We assessed the association between GMV changes (t2-t1) and memory retention in the (**a**) left and (**b**) right JLC using Pearsons’ correlation and observed no significant association in the left JLC (*r* = .27, 95% *CI* = [-0.06, 0.55], *p* = .10, *n* = 37) or right JLC (*r* = .28, 95% *CI* = [-0.04, 0.56], *p* = .09, *n* = 37). When comparing memory retention after splitting the data by median GMV change (t1-t0) in the left **(c,** above: *M* = -5.66, *SD* = 6.91, *n* = 19; below: *M* = -8.89, *SD* = 10.19, *n* = 18) and right JLC **(d,** above: *M* = -6.18, *SD* = 5.80, *n* = 19; below: *M* = -8.33, *SD* = 11.04, *n* = 18), we observed similarly no significant differences for either the left (*t*(35) = -1.13, *d* = -0.26, *p* = .26) or right JLC (*t*(35) = -0.75, *d* = -0.17, *p* = .46). Individual data points are shown over the violin plots. GMV, gray matter volume; R4, fourth retrieval; R5, fifth retrieval; PC, parietal cortex; t0, baseline; t1, 2 h post-learning. Gray, shaded regions indicate the 95 % confidence interval; Bonferroni-corrected; n.s. not significant.

**Fig. S7.**
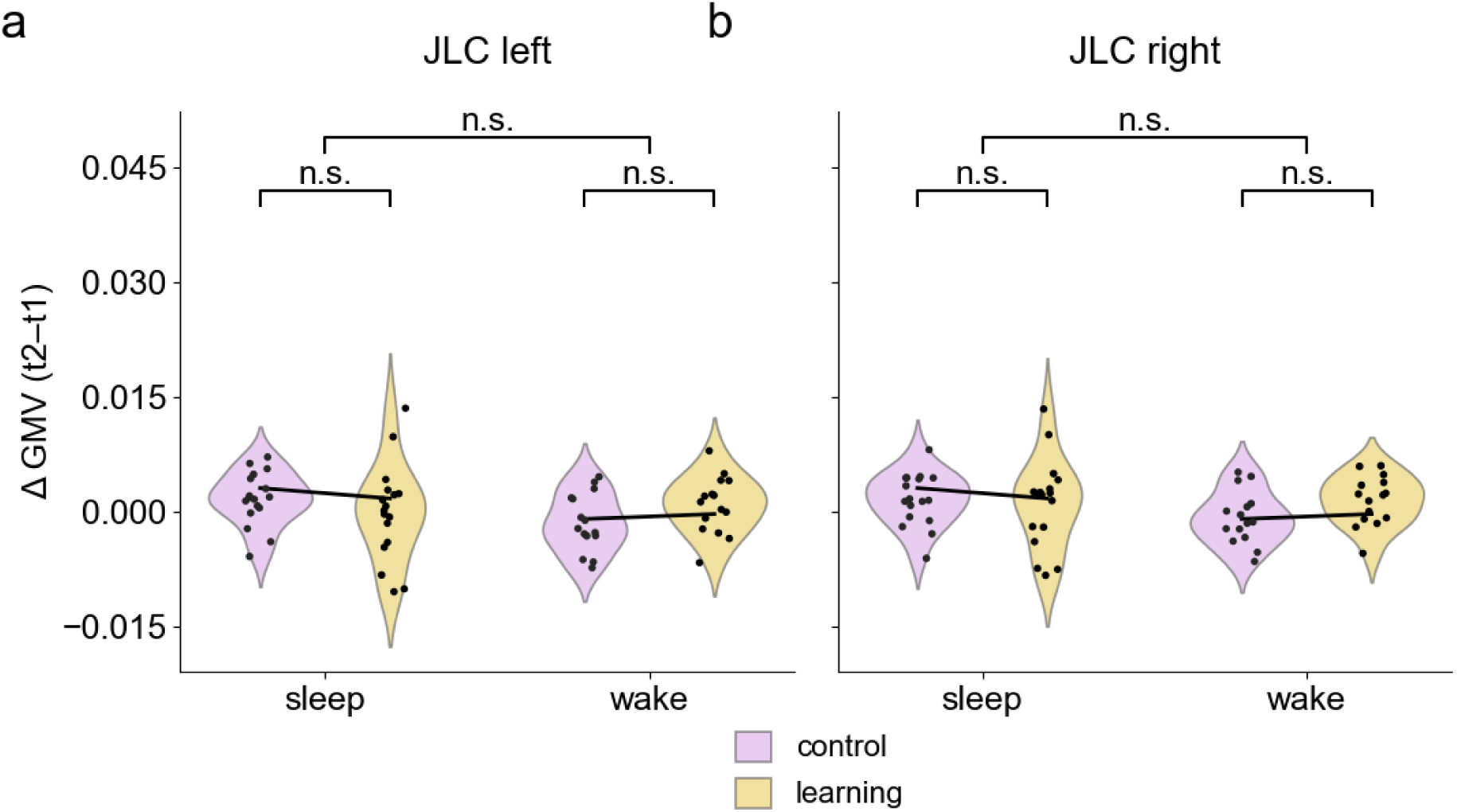
Control analysis analogous to Fig. 5. using juxtapositional lobule cortex (JLC). GMV change between baseline and 12 h post-learning (t2–t1) in the left (**a**) and right (**b**) JLC, stratified by sleep condition (sleep group: studies 1, *n* = 17; wake group: study 1, *n* = 15). The sleep group showed no change in GMV renormalization compared to the no-sleep group in both the left and right JLC (*p* > .05). A 2 × 2 mixed ANOVA found neither the main effect of sleep (left: *F*(1, 30) = 0.80, *p* = .38, η² = .036, right: *F*(1, 30) = 0.78, *p* = .38, η² = .018) or condition (left: *F*(1, 30) = 0.005, *p* = .94, η² = .0001, right: *F*(1, 30) = 0.56, *p* = .46, η² = .018) nor the sleep × condition interaction (left: *F*(1, 30) = 3.66, *p* = .07, η² = .11, right: *F*(1, 30) = 1.98, *p* = .17, η² = .062) to be significant. GMV, gray matter volume; t0, baseline pre-learning; t2, 12 h post-learning; Bonferroni-corrected; n.s. not significant.

## References

Allen, G., Buxton, R. B., Wong, E. C., & Courchesne, E. (1997). Attentional activation of the cerebellum independent of motor involvement. Science, 275(5308), 1940–1943. 10.1126/SCIENCE.275.5308.1940/ASSET/658CDB4D-F6A5-4554-AC24-664F09D94469/ASSETS/GRAPHIC/SE1174887004.JPEG

Ashburner, J. (2007). A fast diffeomorphic image registration algorithm. Neuroimage. https://www.sciencedirect.com/science/article/pii/S1053811907005848

Barsegyan, A., McGaugh, J. L., & Roozendaal, B. (2014). Noradrenergic activation of the basolateral amygdala modulates the consolidation of object-in-context recognition memory. Frontiers in Behavioral Neuroscience, 8(MAY), 89693. 10.3389/FNBEH.2014.00160/TEXT

Bencze, D., Marián, M., Szollosi, Á., Pajkossy, P., Nemecz, Z., Keresztes, A., Hermann, P., Vidnyánszky, Z., & Racsmány, M. (2024). Contribution of the lateral occipital and parahippocampal cortices to pattern separation of objects and contexts. Cerebral Cortex (New York, N.Y.⍰: 1991), 34(7). 10.1093/CERCOR/BHAE295

Bernardi, G., Cecchetti, L., Siclari, F., Buchmann, A., Yu, X., Handjaras, G., Bellesi, M., Ricciardi, E., Kecskemeti, S. R., Riedner, B. A., Alexander, A. L., Benca, R. M., Ghilardi, M. F., Pietrini, P., Cirelli, C., & Tononi, G. (2016). Sleep reverts changes in human gray and white matter caused by wake-dependent training. NeuroImage, 129, 367–377. 10.1016/j.neuroimage.2016.01.020

Bernardinelli, Y., Randall, J., Janett, E., Nikonenko, I., König, S., Jones, E. V., Flores, C. E., Murai, K. K., Bochet, C. G., Holtmaat, A., & Muller, D. (2014). Activity-dependent structural plasticity of perisynaptic astrocytic domains promotes excitatory synapse stability. Current Biology, 24(15), 1679–1688. 10.1016/J.CUB.2014.06.025/ATTACHMENT/C1FF21EB-73F3-4D10-98FF-056351AF275A/MMC5.PDF

Blackwood, N., Ffytche, D., Simmons, A., Bentall, R., Murray, R., & Howard, R. (2004). The cerebellum and decision making under uncertainty. Cognitive Brain Research, 20(1), 46–53. 10.1016/J.COGBRAINRES.2003.12.009

Boven, E., & Cerminara, N. L. (2023). Cerebellar contributions across behavioural timescales: a review from the perspective of cerebro-cerebellar interactions. Frontiers in Systems Neuroscience, 17, 1211530. 10.3389/FNSYS.2023.1211530/TEXT

Boven, E., Pemberton, J., Chadderton, P., Apps, R., & Costa, R. P. (2023). Cerebro-cerebellar networks facilitate learning through feedback decoupling. Nature Communications 2023 14:1, 14(1), 51-. 10.1038/s41467-022-35658-8

Brodt, S., Gais, S., Beck, J., Erb, M., Scheffler, K., & Schönauer, M. (2018). Fast track to the neocortex: A memory engram in the posterior parietal cortex. Science, 362(6418), 1045–1048. 10.1126/SCIENCE.AAU2528/SUPPL_FILE/AAU2528_S1.MP4

Brodt, S., Schönauer, M., Seewald, A., Beck, J., Erb, M., Scheffler, K., & Gais, S. (2023). Memory systems integration in sleep complements rapid systems consolidation in wakefulness. BioRxiv, 2023.03.09.531360. 10.1101/2023.03.09.531360

Broessner, G., Ellerbrock, I., Menz, M. M., Frank, F., Verius, M., Gaser, C., & May, A. (2021). Repetitive T1 Imaging Influences Gray Matter Volume Estimations in Structural Brain Imaging. Frontiers in Neurology, 12, 755749. 10.3389/FNEUR.2021.755749/TEXT

Burton, H., Sinclair, R. J., Wingert, J. R., & Dierker, D. L. (2008). Multiple parietal operculum subdivisions in humans: Tactile activation maps. Somatosensory & Motor Research, 25(3), 149–162. 10.1080/08990220802249275

Bzdok, D., Langner, R., Schilbach, L., Jakobs, O., Roski, C., Caspers, S., Laird, A. R., Fox, P. T., Zilles, K., & Eickhoff, S. B. (2013). Characterization of the temporo-parietal junction by combining data-driven parcellation, complementary connectivity analyses, and functional decoding. NeuroImage, 81, 381–392. 10.1016/J.NEUROIMAGE.2013.05.046

Cairo, T. A., Liddle, P. F., Woodward, T. S., & Ngan, E. T. C. (2004). The influence of working memory load on phase specific patterns of cortical activity. Cognitive Brain Research, 21(3), 377–387. 10.1016/J.COGBRAINRES.2004.06.014

Carter, R. M. K., & Huettel, S. A. (2013). A nexus model of the temporal–parietal junction. Trends in Cognitive Sciences, 17(7), 328–336. 10.1016/J.TICS.2013.05.007

Chen, S. H. A., & Desmond, J. E. (2005). Cerebrocerebellar networks during articulatory rehearsal and verbal working memory tasks. NeuroImage, 24(2), 332–338. 10.1016/J.NEUROIMAGE.2004.08.032

Ciaramelli, E., Grady, C. L., & Moscovitch, M. (2008). Top-down and bottom-up attention to memory: A hypothesis (AtoM) on the role of the posterior parietal cortex in memory retrieval. Neuropsychologia, 46(7), 1828–1851. 10.1016/J.NEUROPSYCHOLOGIA.2008.03.022

Corbetta, M., & Shulman, G. L. (2002). Control of goal-directed and stimulus-driven attention in the brain. Nature Reviews Neuroscience 2002 3:3, 3(3), 201–215. 10.1038/nrn755

Daniels, C., Witt, K., Wolff, S., Jansen, O., & Deuschl, G. (2003). Rate dependency of the human cortical network subserving executive functions during generation of random number series – a functional magnetic resonance imaging study. Neuroscience Letters, 345(1), 25–28. 10.1016/S0304-3940(03)00496-8

Degl’Innocenti, E., & Dell’Anno, M. T. (2023). Human and mouse cortical astrocytes: a comparative view from development to morphological and functional characterization. Frontiers in Neuroanatomy, 17, 1130729. 10.3389/FNANA.2023.1130729

Detre, J. A., Rao, H., Wang, D. J. J., Chen, Y. F., & Wang, Z. (2012). Applications of Arterial Spin Labeled MRI in the Brain. Journal of Magnetic Resonance Imaging, 35(5), 1026. 10.1002/JMRI.23581

Dice, L. R. (1945). Measures of the Amount of Ecologic Association Between Species. Ecology, 26(3), 297–302. 10.2307/1932409

Doniger, G. M., Foxe, J. J., Murray, M. M., Higgins, B. A., Snodgrass, J. G., Schroeder, C. E., & Javitt, D. C. (2000). Activation Timecourse of Ventral Visual Stream Object-recognition Areas: High Density Electrical Mapping of Perceptual Closure Processes. Journal of Cognitive Neuroscience, 12(4), 615– 621. 10.1162/089892900562372

Dordevic, M., Taubert, M., Müller, P., Kaufmann, J., Hökelmann, A., & Müller, N. G. (2018). Brain Gray Matter Volume Is Modulated by Visual Input and Overall Learning Success but Not by Time Spent on Learning a Complex Balancing Task. Journal of Clinical Medicine 2019, Vol. 8, Page 9, 8(1), 9. 10.3390/JCM8010009

Doricchi, F. (2022). The functions of the temporal–parietal junction. Handbook of Clinical Neurology, 187, 161–177. 10.1016/B978-0-12-823493-8.00020-1

Draganski, B., Gaser, C., Busch, V., Schuierer, G., Bogdahn, U., & May, A. (2004). Changes in grey matter induced by training. Nature 2004 427:6972, 427(6972), 311–312. 10.1038/427311a

Draganski, B., Gaser, C., Kempermann, G., Kuhn, H. G., Winkler, J., Büchel, C., & May, A. (2006). Temporal and Spatial Dynamics of Brain Structure Changes during Extensive Learning. Journal of Neuroscience, 26(23), 6314–6317. 10.1523/JNEUROSCI.4628-05.2006

Eriksson, P. S., Perfilieva, E., Björk-Eriksson, T., Alborn, A. M., Nordborg, C., Peterson, D. A., & Gage, F. H. (1998). Neurogenesis in the adult human hippocampus. Nature Medicine 1998 4:11, 4(11), 1313–1317. 10.1038/3305

Folvik, L., Sneve, M. H., Ness, H. T., Vidal-Piñeiro, D., Raud, L., Geier, O. M., Walhovd, K. B., & Fjell, A. M. (2023). Sustained upregulation of widespread hippocampal–neocortical coupling following memory encoding. Cerebral Cortex, 33(8), 4844–4858. 10.1093/CERCOR/BHAC384

Forstmeier, W., & Schielzeth, H. (2011). Cryptic multiple hypotheses testing in linear models: Overestimated effect sizes and the winner’s curse. Behavioral Ecology and Sociobiology, 65(1), 47–55. 10.1007/S00265-010-1038-5/FIGURES/3

Frankland, P. W., & Bontempi, B. (2005). The organization of recent and remote memories. Nature Reviews Neuroscience 2005 6:2, 6(2), 119–130. 10.1038/nrn1607

Friedman, N., Malovani, C., Perets, I., Kenin, E., Bernstein-Eliav, M., & Tavor, I. (2025). Continuous Diffusion-Detected Neuroplasticity during Motor Learning. Journal of Neuroscience, 45(24). 10.1523/JNEUROSCI.1152-24.2025

Friston, K. J. (2003). Statistical Parametric Mapping. Neuroscience Databases, 237–250. 10.1007/978-1-4615-1079-6_16

Fromm, A. E., Abdelmotaleb, M., Olschewski, F., Limanowski, J., Meinzer, M., Flöel, A., & Antonenko, D. (2026). Brain Regions Involved in Object-Location Memory Across the Human Lifespan: A Systematic Review and Activation Likelihood Estimation Meta-Analysis of Task-Based fMRI. BioRxiv, 2026.07.01.735849. 10.64898/2026.07.01.735849

Gaser, C., Dahnke, R., Thompson, P. M., Kurth, F., & Luders, E. (2024). CAT: a computational anatomy toolbox for the analysis of structural MRI data. GigaScience, 13, 1–13. 10.1093/GIGASCIENCE/GIAE049

Gillis, M. M., Garcia, S., & Hampstead, B. M. (2016). Working memory contributes to the encoding of object location associations: Support for a 3-part model of object location memory. Behavioural Brain Research, 311, 192–200. 10.1016/J.BBR.2016.05.037

Gorgolewski, K. J., Varoquaux, G., Rivera, G., Schwarz, Y., Ghosh, S. S., Maumet, C., Sochat, V. V., Nichols, T. E., Poldrack, R. A., Poline, J. B., Yarkoni, T., & Margulies, D. S. (2015). NeuroVault.org: a web-based repository for collecting and sharing unthresholded statistical maps of the human brain. Frontiers in Neuroinformatics, 9(APR). 10.3389/FNINF.2015.00008

Grill-Spector, K., Kushnir, T., Edelman, S., Avidan, G., Itzchak, Y., & Malach, R. (1999). Differential processing of objects under various viewing conditions in the human lateral occipital complex. Neuron, 24(1), 187–203. 10.1016/S0896-6273(00)80832-6/ASSET/9EC9E66A-0F55-41FE-9C53-84E6FC9A8761/MAIN.ASSETS/GR8.JPG

Grill-Spector, K., Kushnir, T., Hendler, T., Edelman, S., Itzchak, Y., & Malach, R. (1998). A Sequence of Object-Processing Stages Revealed by fMRI in the Human Occipital Lobe. Hum. Brain Mapping, 6, 316–328. 10.1002/(SICI)1097-0193(1998)6:4

Guerra-Carrillo, B., MacKey, A. P., & Bunge, S. A. (2014). Resting-state fMRI: A window into human brain plasticity. Neuroscientist, 20(5), 522–533. 10.1177/1073858414524442/ASSET/D02ED20F-DA37-45CF-89B2-7A79C982FB4F/ASSETS/IMAGES/LARGE/10.1177_1073858414524442-FIG4.JPG

Hampstead, B. M., Stringer, A. Y., Stilla, R. F., Amaraneni, A., & Sathian, K. (2011). Where did I put that? Patients with amnestic mild cognitive impairment demonstrate widespread reductions in activity during the encoding of ecologically relevant object-location associations. Neuropsychologia, 49(9), 2349–2361. 10.1016/J.NEUROPSYCHOLOGIA.2011.04.008

Harrington, D. L., Boyd, L. A., Mayer, A. R., Sheltraw, D. M., Lee, R. R., Huang, M., & Rao, S. M. (2004). Neural representation of interval encoding and decision making. Cognitive Brain Research, 21(2), 193–205. 10.1016/J.COGBRAINRES.2004.01.010

Himmer, L., Schönauer, M., Heib, D. P. J., Schabus, M., & Gais, S. (2019). Rehearsal initiates systems memory consolidation, sleep makes it last. Science Advances, 5(4). 10.1126/SCIADV.AAV1695/SUPPL_FILE/AAV1695_SM.PDF

Jiang, P., Chen, C., Liu, X. B., Pleasure, D. E., Liu, Y., & Deng, W. (2016). Human iPSC-Derived Immature Astroglia Promote Oligodendrogenesis by Increasing TIMP-1 Secretion. Cell Reports, 15(6), 1303– 1315. 10.1016/J.CELREP.2016.04.011/ATTACHMENT/6B41B430-9076-4BA7-8B1A-E2A2ED829EA6/MMC2.PDF

Johnson, W. E., Li, C., & Rabinovic, A. (2007). Adjusting batch effects in microarray expression data using empirical Bayes methods. Biostatistics, 8(1), 118–127. 10.1093/BIOSTATISTICS/KXJ037

Jung, P., Baumgärtner, U., Stoeter, P., & Treede, R. D. (2009). Structural and functional asymmetry in the human parietal opercular cortex. Journal of Neurophysiology, 101(6), 3246–3257. 10.1152/JN.91264.2008/ASSET/IMAGES/LARGE/Z9K0060995020005.JPEG

Keifer, O. P., Hurt, R. C., Gutman, D. A., Keilholz, S. D., Gourley, S. L., & Ressler, K. J. (2015). Voxel-based morphometry predicts shifts in dendritic spine density and morphology with auditory fear conditioning. Nature Communications 2015 6:1, 6(1), 7582-. 10.1038/ncomms8582

Kempermann, G., Song, H., & Gage, F. H. (2015). Neurogenesis in the Adult Hippocampus. Cold Spring Harbor Perspectives in Biology, 7(9), a018812. 10.1101/CSHPERSPECT.A018812

Kostadinov, D., Beau, M., Pozo, M. B., & Häusser, M. (2019). Predictive and reactive reward signals conveyed by climbing fiber inputs to cerebellar Purkinje cells. Nature Neuroscience 2019 22:6, 22(6), 950–962. 10.1038/s41593-019-0381-8

Kourtzi, Z., Betts, L. R., Sarkheil, P., & Welchman, A. E. (2005). Distributed Neural Plasticity for Shape Learning in the Human Visual Cortex. PLOS Biology, 3(7), e204. 10.1371/JOURNAL.PBIO.0030204

Kuhn, H. G., Palmer, T. D., & Fuchs, E. (2001). Adult neurogenesis: A compensatory mechanism for neuronal damage. European Archives of Psychiatry and Clinical Neuroscience, 251(4), 152–158. 10.1007/S004060170035/METRICS

Kumral, D., Lenders, A., & Schönauer, M. (2026). Learning sculpts microstructure in real time: evidence from dense temporal sampling of diffusion MRI. BioRxiv, 2026.06.24.734293. 10.64898/2026.06.24.734293

Lanjewar, S. N., & Sloan, S. A. (2021). Growing Glia: Cultivating Human Stem Cell Models of Gliogenesis in Health and Disease. Frontiers in Cell and Developmental Biology, 9, 649538. 10.3389/FCELL.2021.649538/XML

Lewis, L. D., Setsompop, K., Rosen, B. R., & Polimeni, J. R. (2018). Stimulus-dependent hemodynamic response timing across the human subcortical-cortical visual pathway identified through high spatiotemporal resolution 7T fMRI. NeuroImage, 181, 279. 10.1016/J.NEUROIMAGE.2018.06.056

Liu, Y. J., Spangenberg, E. E., Tang, B., Holmes, T. C., Green, K. N., & Xu, X. (2021). Microglia Elimination Increases Neural Circuit Connectivity and Activity in Adult Mouse Cortex. Journal of Neuroscience, 41(6), 1274–1287. 10.1523/JNEUROSCI.2140-20.2020

Lövdén, M., Wenger, E., Mårtensson, J., Lindenberger, U., & Bäckman, L. (2013). Structural brain plasticity in adult learning and development. Neuroscience & Biobehavioral Reviews, 37(9), 2296–2310. 10.1016/J.NEUBIOREV.2013.02.014

Månsson, K. N. T., Cortes, Di. S., Manzouri, A., Li, T. Q., Hau, S., & Fischer, H. (2020). Viewing Pictures Triggers Rapid Morphological Enlargement in the Human Visual Cortex. Cerebral Cortex, 30(3), 851– 857. 10.1093/CERCOR/BHZ131

Maule, F., Barchiesi, G., Brochier, T., & Cattaneo, L. (2015). Haptic Working Memory for Grasping: the Role of the Parietal Operculum. Cerebral Cortex, 25(2), 528–537. 10.1093/CERCOR/BHT252

McGaugh, J. L. (2000). Memory - A century of consolidation. Science, 287(5451), 248–251. 10.1126/SCIENCE.287.5451.248

McGaugh, J. L. (2004). The amygdala modulates the consolidation of memories of emotionally arousing experiences. Annual Review of Neuroscience, 27(Volume 27, 2004), 1–28. 10.1146/ANNUREV.NEURO.27.070203.144157/CITE/REFWORKS

Moore, M. J., Jenkinson, M., Griffanti, L., Huygelier, H., Gillebert, C. R., & Demeyere, N. (2023). A comparison of lesion mapping analyses based on CT versus MR imaging in stroke. Neuropsychologia, 184, 108564. 10.1016/J.NEUROPSYCHOLOGIA.2023.108564

Naegel, S., Hagenacker, T., Theysohn, N., Diener, H.-C., Katsarava, Z., Obermann, M., & Holle, D. (2017). Short Latency Gray Matter Changes in Voxel-Based Morphometry following High Frequent Visual Stimulation. Wiley Online Library, 2017. 10.1155/2017/1397801

Nakamura, K., Brown, R. A., Narayanan, S., Louis Collins, D., & Arnold, D. L. (2015). Diurnal fluctuations in brain volume: Statistical analyses of MRI from large populations ⋆. 10.1016/j.neuroimage.2015.05.077

Nierhaus, T., Vidaurre, C., Sannelli, C., Mueller, K. R., & Villringer, A. (2021). Immediate brain plasticity after one hour of brain–computer interface (BCI). Journal of Physiology, 599(9), 2435–2451. 10.1113/JP278118

Olivo, G., Lövdén, M., Manzouri, A., Terlau, L., Jenner, B., Jafari, A., Petersson, S., Li, T. Q., Fischer, H., & Månsson, K. N. T. (2022). Estimated gray matter volume rapidly changes after a short motor task. Cerebral Cortex, 32(19), 4356–4369. 10.1093/CERCOR/BHAB488

Pan, Y., & Monje, M. (2020). Activity Shapes Neural Circuit Form and Function: A Historical Perspective. Journal of Neuroscience, 40(5), 944–954. 10.1523/JNEUROSCI.0740-19.2019

Parkhurst, C. N., Yang, G., Ninan, I., Savas, J. N., Yates, J. R., Lafaille, J. J., Hempstead, B. L., Littman, D. R., & Gan, W. B. (2013). Microglia promote learning-dependent synapse formation through brain-derived neurotrophic factor. Cell, 155(7), 1596–1609. 10.1016/J.CELL.2013.11.030/ATTACHMENT/736C3008-D474-4450-80B7-9A47312AD476/MMC2.PDF

Patel, G. H., Sestieri, C., & Corbetta, M. (2019). The evolution of the temporoparietal junction and posterior superior temporal sulcus. Cortex, 118, 38–50. 10.1016/J.CORTEX.2019.01.026

Pereira, A. C., Huddleston, D. E., Brickman, A. M., Sosunov, A. A., Hen, R., McKhann, G. M., Sloan, R., Gage, F. H., Brown, T. R., & Small, S. A. (2007). An in vivo correlate of exercise-induced neurogenesis in the adult dentate gyrus. Proceedings of the National Academy of Sciences of the United States of America, 104(13), 5638–5643. 10.1073/PNAS.0611721104/SUPPL_FILE/11721FIG5.PDF

Phelps, E. A. (2004). Human emotion and memory: interactions of the amygdala and hippocampal complex. Current Opinion in Neurobiology, 14(2), 198–202. 10.1016/J.CONB.2004.03.015

Ponti, G., Obernier, K., & Alvarez-Buylla, A. (2013). Cell Cycle Lineage progression from stem cells to new neurons in the adult brain ventricular-subventricular zone. 10.4161/cc.24984

Postma, A., Kessels, R. P. C., & van Asselen, M. (2008). How the brain remembers and forgets where things are: The neurocognition of object–location memory. Neuroscience & Biobehavioral Reviews, 32(8), 1339–1345. 10.1016/J.NEUBIOREV.2008.05.001

Rhyu, I. J., Bytheway, J. A., Kohler, S. J., Lange, H., Lee, K. J., Boklewski, J., McCormick, K., Williams, N. I., Stanton, G. B., Greenough, W. T., & Cameron, J. L. (2010). Effects of aerobic exercise training on cognitive function and cortical vascularity in monkeys. Neuroscience, 167(4), 1239–1248. 10.1016/J.NEUROSCIENCE.2010.03.003

Roozendaal, B., Castello, N. A., Vedana, G., Barsegyan, A., & McGaugh, J. L. (2008). Noradrenergic activation of the basolateral amygdala modulates consolidation of object recognition memory. Neurobiology of Learning and Memory, 90(3), 576–579. 10.1016/J.NLM.2008.06.010

Ross, R. S., & Slotnick, S. D. (2008). The Hippocampus is Preferentially Associated with Memory for Spatial Context. Journal of Cognitive Neuroscience, 20(3), 432–446. 10.1162/JOCN.2008.20035

Sagi, Y., Tavor, I., Hofstetter, S., Tzur-Moryosef, S., Blumenfeld-Katzir, T., & Assaf, Y. (2012). Learning in the Fast Lane: New Insights into Neuroplasticity. Neuron, 73(6), 1195–1203. 10.1016/j.neuron.2012.01.025

Schmidt, S., Gull, S., Herrmann, K. H., Boehme, M., Irintchev, A., Urbach, A., Reichenbach, J. R., Klingner, C. M., Gaser, C., & Witte, O. W. (2021). Experience-dependent structural plasticity in the adult brain: How the learning brain grows. NeuroImage, 225, 117502. 10.1016/J.NEUROIMAGE.2020.117502

Shen, X., Tokoglu, F., Papademetris, X., & Constable, R. T. (2013). Groupwise whole-brain parcellation from resting-state fMRI data for network node identification. NeuroImage, 82, 403–415. 10.1016/J.NEUROIMAGE.2013.05.081

Sirigu, A., & Desmurget, M. (2021). Somatosensory awareness in the parietal operculum. Brain, 144(12), 3558–3560. 10.1093/BRAIN/AWAB415

Sokolov, A. A., Miall, R. C., & Ivry, R. B. (2017). The Cerebellum: Adaptive Prediction for Movement and Cognition. Trends in Cognitive Sciences, 21(5), 313–332. 10.1016/J.TICS.2017.02.005/ASSET/335B493A-4574-43DC-AA40-007D1EF6224F/MAIN.ASSETS/GR2.JPG

Stoodley, C. J., & Schmahmann, J. D. (2009). Functional topography in the human cerebellum: A meta-analysis of neuroimaging studies. NeuroImage, 44(2), 489–501. 10.1016/J.NEUROIMAGE.2008.08.039

Takashima, A., Nieuwenhuis, I. L. C., Jensen, O., Talamini, L. M., Rijpkema, M., & Fernández, G. (2009). Shift from Hippocampal to Neocortical Centered Retrieval Network with Consolidation. Journal of Neuroscience, 29(32), 10087–10093. 10.1523/JNEUROSCI.0799-09.2009

Tardif, C. L., Steele, C. J., Lampe, L., Bazin, P. L., Ragert, P., Villringer, A., & Gauthier, C. J. (2017). Investigation of the confounding effects of vasculature and metabolism on computational anatomy studies. NeuroImage, 149, 233–243. 10.1016/J.NEUROIMAGE.2017.01.025

Taubert, M., Draganski, B., Anwander, A., Müller, K., Horstmann, A., Villringer, A., & Ragert, P. (2010). Dynamic Properties of Human Brain Structure: Learning-Related Changes in Cortical Areas and Associated Fiber Connections. Journal of Neuroscience, 30(35), 11670–11677. 10.1523/JNEUROSCI.2567-10.2010

Taubert, M., Mehnert, J., Pleger, B., & Villringer, A. (2016). Rapid and specific gray matter changes in M1 induced by balance training. NeuroImage, 133, 399–407. 10.1016/J.NEUROIMAGE.2016.03.017

Theodosis, D. T., Poulain, D. A., & Oliet, S. H. R. (2008). Activity-dependent structural and functional plasticity of astrocyte-neuron interactions. Physiological Reviews, 88(3), 983–1008. 10.1152/PHYSREV.00036.2007/ASSET/IMAGES/LARGE/Z9J0030824810009.JPEG

Tononi, G., & Cirelli, C. (2006). Sleep function and synaptic homeostasis. Sleep Medicine Reviews, 10(1), 49–62. 10.1016/J.SMRV.2005.05.002

Trefler, A., Sadeghi, N., Thomas, A. G., Pierpaoli, C., Baker, C. I., & Thomas, C. (2016). Impact of time-of-day on brain morphometric measures derived from T1-weighted magnetic resonance imaging. NeuroImage, 133, 41–52. 10.1016/J.NEUROIMAGE.2016.02.034

Tremblay, M. Ě., Lowery, R. L., & Majewska, A. K. (2010). Microglial Interactions with Synapses Are Modulated by Visual Experience. PLOS Biology, 8(11), e1000527. 10.1371/JOURNAL.PBIO.1000527

Uhlig, M., Reinelt, J. D., Lauckner, M. E., Kumral, D., Schaare, H. L., Mildner, T., Babayan, A., Möller, H. E., Engert, V., Villringer, A., & Gaebler, M. (2023). Rapid volumetric brain changes after acute psychosocial stress. NeuroImage, 265, 119760. 10.1016/J.NEUROIMAGE.2022.119760

Verra, L., Renz, F., & Schuck, N. (2024). Neuroscience Methods for Investigating Brain Plasticity (Chapter to appear in The Oxford Handbook of Cognitive Enhancement and Brain Plasticity). https://europepmc.org/article/ppr/ppr951783

Villa, S., Kleinschroth, F., Schönauer, M., & Kumral, D. (2026). Experience-dependent rapid structural changes in the human brain: A systematic review. Neuroscience & Biobehavioral Reviews, 187, 106736. 10.1016/J.NEUBIOREV.2026.106736

Wenger, E., Brozzoli, C., Lindenberger, U., & Lövdén, M. (2017). Expansion and Renormalization of Human Brain Structure During Skill Acquisition. Trends in Cognitive Sciences, 21(12), 930–939. 10.1016/J.TICS.2017.09.008/ASSET/ADDC83A5-E1BE-4DED-A092-5D5B0655EDEB/MAIN.ASSETS/GR1.JPG

Westerlund, U., Moe, M. C., Varghese, M., Berg-Johnsen, J., Ohlsson, M., Langmoen, I. A., & Svensson, M. (2003). Stem cells from the adult human brain develop into functional neurons in culture. Experimental Cell Research, 289(2), 378–383. 10.1016/S0014-4827(03)00291-X

Wolpert, D. M., Miall, R. C., & Kawato, M. (1998). Internal models in the cerebellum. Trends in Cognitive Sciences, 2(9), 338–347. 10.1016/S1364-6613(98)01221-2

Zaretskaya, N., Fink, E., Arsenovic, A., & Ischebeck, A. (2023). Fast and functionally specific cortical thickness changes induced by visual stimulation. Cerebral Cortex, 33(6), 2823–2837. 10.1093/CERCOR/BHAC244

Zatorre, R. J., Fields, R. D., & Johansen-Berg, H. (2012). Plasticity in gray and white: neuroimaging changes in brain structure during learning. Nature Neuroscience 2012 15:4, 15(4), 528–536. 10.1038/nn.3045

